# Assembly of the *Drosophila* mushroom body circuit and its regulation by Semaphorin 1a

**DOI:** 10.1101/835595

**Authors:** Chen-Han Lin, Suewei Lin

## Abstract

The *Drosophila* mushroom body (MB) is a learning and memory center in the fly brain. It is the most extensively studied brain structure in insects, but we know little about the molecular and cellular mechanisms underlying assembly of its neural circuit. The MB is composed of around 2200 intrinsic Kenyon cells (KCs), whose axons are bundled to form multiple MB lobes. The MB lobes are innervated by a large number of extrinsic neurons. Twenty types of dopaminergic neurons (DANs) and 21 types of MB output neurons (MBONs) have been identified. Each type of these extrinsic neurons innervates specific compartments or zones in the MB lobes. Here, we characterize the assembly of the MB circuit and reveal several intriguing features of the process. The DANs and MBONs innervate zones in the MB vertical lobes in specific sequential orders. Innervation of DAN axons in some zones precedes that of MBON dendrites, and *vice versa* in other zones. MBON and DAN innervations are largely independent of each other. Removing one type of extrinsic neuron during early development has a limited effect on the MB lobe innervations of the other type of extrinsic neurons. However, KC axons are essential for zonal elaboration of DAN axons and MBON dendrites. Competition also exists between MB zones for some MBONs, so when the cognate zones for these MBONs are missing, their dendrites are misdirected to other zones. Finally, we identify Semaphorin 1a (Sema1a) as a crucial guidance molecule for MBON dendrites to innervate specific MB lobe zones. Ectopic expression of Sema1a in some DANs is sufficient to re-direct their dendrites to those zones, demonstrating a potential to rewire the MB circuit. Taken together, our work provides an initial characterization of the cellular and molecular mechanisms underlying MB circuit assembly.

## Introduction

During brain development, large numbers of neurons are precisely assembled in three-dimensional space to form functional networks that ultimately give rise to complex behaviors. The assembly process is regulated by intricate cellular and molecular mechanisms. Disruption of these mechanisms can lead to severe behavioral disorders or even death. Since Roger Sperry proposed his famous chemoaffinity hypothesis (Sperry, 1963), postulating that the precise wiring of the nervous system is established by specific chemical affinities generated from individual neurons, a large number of guidance molecules and principles governing neural circuit development has been discovered (Chédotal and Richards, 2010; Dickson and Zou, 2010; Hong and Luo, 2014; Kolodkin and Tessier-Lavigne, 2011; Sanes and Yamagata, 2009). However, our understanding of neural circuit assembly remains profoundly insufficient and we still cannot explain fully the mechanisms underpinning the wiring process of any neural circuit. Furthermore, given the remarkably diverse cell types and circuit configurations in the brain, additional guiding strategies and molecules surely await discovery.

The genetically tractable yet sufficiently complex nervous system of the fruit fly *Drosophila* has proven to be an excellent model for studying neural development. Many important principles of molecular guidance and neural circuit organization were first discovered in the fly (Evans, 2016; Hong and Luo, 2014; Kolodkin and Tessier-Lavigne, 2011; Plazaola-Sasieta et al., 2017; Tessier-Lavigne and Goodman, 1996). Most fly studies have focused on the ventral nerve cord, neuromuscular systems, antennal lobes, and optical lobes, but assembly of the central brain circuits has only begun to be explored (Oliva et al., 2016; Xie et al., 2017, 2019).

The *Drosophila* mushroom body (MB) is a multi-lobe structure at the center of the fly brain. It is a computational center mediating learning and memory (de Belle and Heisenberg, 1994; Heisenberg, 2003), sleep (Joiner et al., 2006; Pitman et al., 2006), and motivated behavior (Krashes et al., 2009; Senapati et al., 2019; Tsao et al., 2018). The MB circuit is composed of three major types of neurons—Kenyon cells (KCs), MB output neurons (MBONs), and dopaminergic neurons (DANs) (Aso et al., 2014). There are around 2200 KCs in each fly brain hemisphere. They are the intrinsic MB neurons whose axonal bundles form five MB lobes, with the α and αʹ lobes extending vertically and the β, βʹ and γ lobes projecting horizontally. The MB lobes are innervated by the dendrites of ∼34 MBONs and the axons of ∼130 DANs. Based on their morphology and molecular markers, 21 types of MBONs and 20 types of DANs have been identified. Each type of MBON or DAN extends its dendrites or axons, respectively, into specific compartments or zones of the MB lobes. Together, MBONs and DANs subdivide the MB lobes into 15 distinct zones. Most MBONs and DANs innervate only one or two zones. Given that the MB lobes are bundles of continuous axonal processes, the way in which each type of MBON and DAN synapses with specific domains of the MB lobes represents an elaborated form of subcellular targeting, through a process that is not yet well understood.

MB development has been studied extensively, but mainly focused on the KCs. KCs can be classified into three major types, i.e., γ, αʹβʹ, and αβ. They are sequentially produced from four neuroblasts during development (Lee et al., 1999). Several molecular mechanisms have been identified as governing the specification of KC cell fates (Liu et al., 2019, 2015; Marchetti and Tavosanis, 2017; Zhu et al., 2006). Targeting and pruning of KC axons have also been well characterized (Awasaki and Ito, 2004; Boulanger et al., 2011; Boyle et al., 2006; Lee et al., 2000; Lin et al., 2009; Ng et al., 2002; Rabinovich et al., 2016; Shimizu et al., 2011; Shin and DiAntonio, 2011; Smith-Trunova et al., 2015; Wang et al., 2002; Yu et al., 2013; Zwarts et al., 2016). In contrast, much less is known about the development of MBONs or DANs and how they are assembled into the MB circuit. To fill this knowledge gap, we first characterized the targeting of MBON dendrites and DAN axons to the MB vertical lobes during the pupal stage when the adult MB circuit is being assembled. We then used ablation experiments to probe the requirement of MBONs, DANs, and KCs for the assembly of the MB circuit. Finally, we identified that *semaphorin 1a* (*sema1a*), a classical guidance molecule (Kolodkin et al., 1993), is expressed in some MBONs, which is critical for their dendritic targeting to specific MB lobe zones. Ectopic expression of *sema1a* in some DANs is sufficient to redirect their dendrites to those zones. Our work provides the first characterization of the assembly of the MB circuit and reveals a role for *sema1a* in the dendritic targeting of MBONs.

## Results

### The innervation timings of DAN axons and MOBN dendrites in the MB vertical lobes

To investigate how the MB circuits are assembled, we first endeavored to identify GAL4 lines in which individual types of DANs and MBONs are labeled throughout their morphogenesis. We focused on the MBONs and DANs that innervate the MB vertical lobes because they are fewer in type and number than those innervating the horizontal lobes (Aso et al., 2014). Among all of the GAL4 lines we assessed (**Table S1**), *MB058B-splitGAL4* and *MB091C-splitGAL4* specifically label PPL1-αʹ2α2 DANs and MBON-αʹ2, respectively, throughout their development (**Fig. 1A-L’**). Since PPL1-αʹ2α2 DANs and MBON-αʹ2 are the only DANs and MBONs that innervate the αʹ2 zone (Aso et al., 2014), we could study the innervation order of DAN axons and MBON dendrites in the same MB lobe zone by following the development of these neurons. We used *MB058B* and *MB091C* to drive the expression of mCD8::GFP in PPL1-αʹ2α2 DANs and MBON-αʹ2 and checked their morphologies at various times after puparium formation (APF). The PPL1-αʹ2α2 DANs begin to touch the MB lobes at 18 h APF, starting from the lateral surface of the αʹ2 zone (**Fig. 1A-A’**). At 24 h APF, innervation became more obvious, with axons extending ventrally along the interfaces between the αʹ and α lobes (**Fig. 1B-B’ and M-M’’**). Dense zonal innervation of the PPL1-αʹ2α2 axons was observed at 36 h APF (**Fig. 1D-D’**). Dendritic targeting of MBON-αʹ2 to the αʹ2 zone occurred at a later time-point. At 42 h APF, MBON-αʹ2 dendrites only began to touch the medial surface of the αʹ2 zone (**Fig. 1K-K’**), and dense zonal elaboration of the MBON-αʹ2 dendrites was only seen at 48 h APF (**Fig. 1L-L’**). Therefore, in the αʹ2 zone, DAN axons innervate the MB lobes earlier than MBON dendrites.

**Figure 1.**
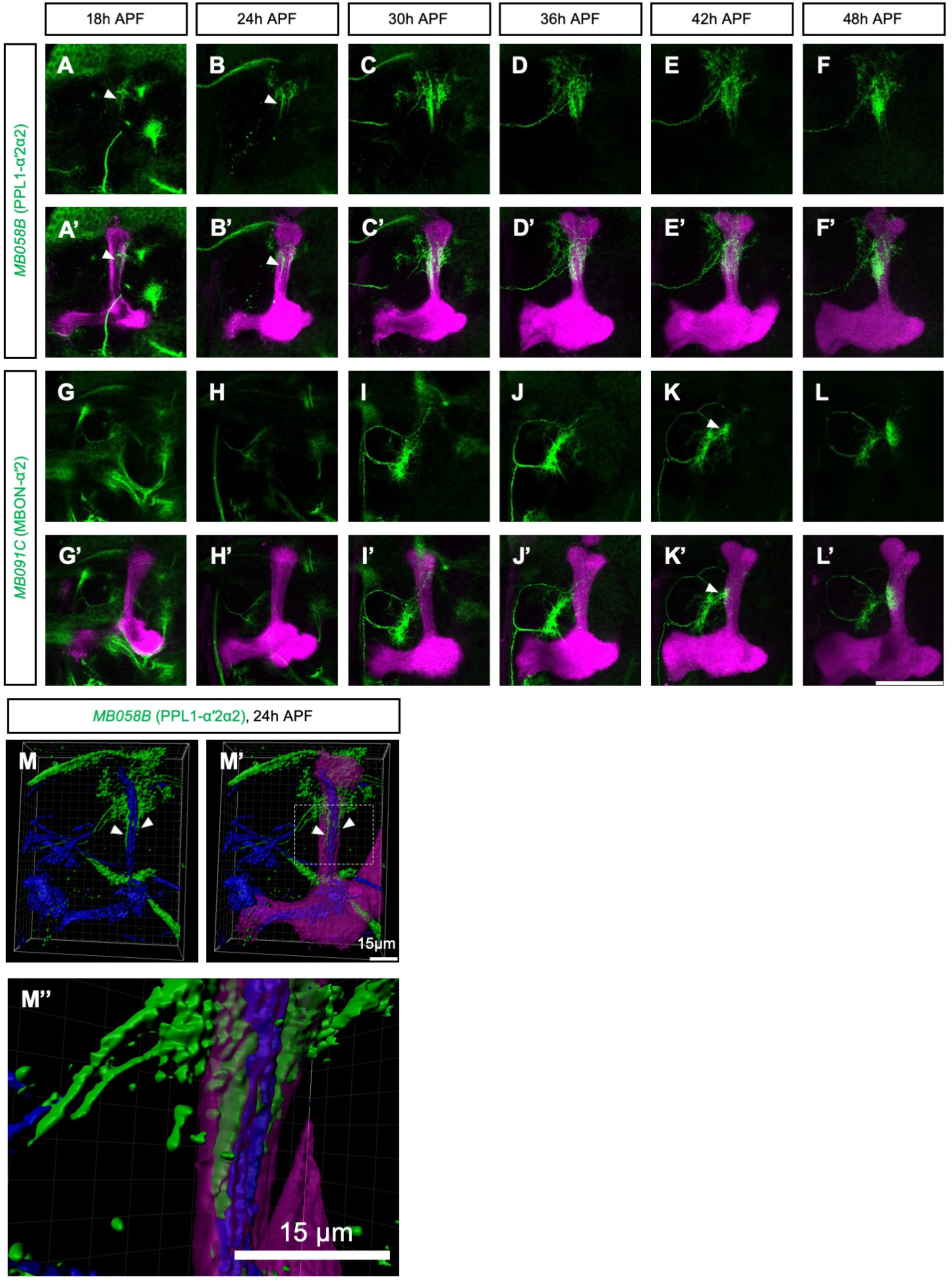
Timing of innervations by DANs and MBONs targeting the αʹ2 zone. **(A-F’)** Axonal innervation in the MB vertical lobes by PPL1-αʹ2α2 DANs, labeled by *MB058B-splitGAL4* (green, A-F’), at different hours after puparium formation (APF). The brains were counterstained with *LexAop-rCD2::RFP* driven by *MB247-LexA::p65* (magenta, A’-F’). The white arrowheads in (A-B’) indicate where the axons start to innervate the MB lobes. **(G-L’)** Dendritic innervation in the MB vertical lobes by MBON-αʹ2, labeled by *MB091C-splitGAL4* (green, G-L’), at different hours APF. The brains were counterstained with *LexAop-rCD2::RFP* driven by *MB247-LexA::p65* (magenta, G’-L’). The white arrowheads in (K-K’) indicate where the dendrites start to innervate the MB lobes. **(M-M’)** 3-D images of the PPL1-αʹ2α2 axons in the MB lobes. Axons were labeled with *UAS-mCD8::GFP* driven by *MB058B-splitGAL4* (green, M-M’) and all MB lobes were labeled with *LexAop-rCD2::RFP* driven by *MB247-LexA::p65* (magenta, M’-M’), and the α and β lobes were labeled with anti-FasII antibody (blue, M-M’). The white arrowheads in indicate the axonal processes extending at the junctions between the α and αʹ lobes. **(M’’)** A magnified view of the area outlined by a white dashed square in (M’). Scale bar in (A-L’): 50 µm. Scale bar in (M-M’’): 15 µm. Detailed genotypes are listed in Table S2.

Interestingly, it seems that DAN axons do not always precede MBON dendrites in all MB lobe zones. For instance, in the αʹ3 zone, MBON-αʹ3 dendrites likely innervate the MB lobes earlier than the PPL1-αʹ3 axons. Innervation by MBON-αʹ3 dendrites (labeled with *MB027B-splitGAL4*) was observed at 6 h APF, i.e., much earlier than observed for neighboring MBON-αʹ2 (**Fig. S1**). However, when we labeled the DANs with antibodies against tyrosine hydroxylase (TH), an enzyme required for the synthesis of dopamine (Neckameyer and Quinn, 1989), we found that TH expression is sequentially turned on, beginning at the base of the vertical lobes and finishing at the tips of the α and αʹ lobes (**Fig. S2**). Expression of TH in the αʹ3 zone only became obvious at 54 h APF, representing the latest time-point among all vertical compartments. Since anti-TH signal was first seen in the αʹ2 and α2 zones at 24 h APF, TH expression appears to correlate with the innervations by DANs (**Fig. 1B-B’ and Fig. S2**). These data suggest that MBON dendrites precede DAN axons in the αʹ3 zone.

### DANs and MBONs independently innervate MB zones

Given that the axons of PPL1-αʹ2α2 DANs precede the MBON-αʹ2 dendrites in the αʹ2 zone, next we examined the possibility that DAN axons provide guidance cues for MBON dendrites. We genetically ablated PPL1-αʹ2α2 DANs by ectopically expressing the apoptotic genes *head involution defective* (*hid*) and *reaper* (*rpr*) (Grether et al., 1995; White et al., 1994) in these neurons using *MB058B-splitGAL4*. No *MB058B-*positive cells were seen in PPL1-αʹ2α2 DAN-ablated adult brains, and the αʹ2 and α2 MB lobe zones lacked anti-TH staining signal (**Fig. 2A-A’ and B-B’**). Ablation of PPL1-αʹ2α2 DANs occurred quite early during development and all *MB058B*-positive neurites were eliminated before 24 h APF, i.e., by the time PPL1-αʹ2α2 DANs start to innervate MB lobes in wild-type brains (**Fig. S3**). TH-negative zones in the PPL1-αʹ2α2 DAN-ablated brains suggest that axon-axon contacts between neighboring DANs have a minimal effect in determining the borders of DAN axon zonal networks in MB lobes. Ablation of PPL1-αʹ2α2 DANs did not affect the dendritic targeting of MBON-αʹ2 (labeled with *R20G03-LexA::P65*-driven *rCD2-RFP*), suggesting that PPL1-αʹ2α2 DANs do not provide guidance cues for MBON-αʹ2 dendrites (**Fig. 2A’’-A’’’ and B’’-B’’’**). Ablation of PPL1-αʹ2α2 DANs also had no effect on the dendritic elaboration of MBON-αʹ3 (labeled with *R53F03-LexA::P65*-driven *rCD2-RFP*) or MBON-γ2αʹ1 (labeled with *R25D01-LexA::P65*-driven *rCD2-RFP*) in the neighboring zones (**Fig. S4**). Similarly, ablation of MBON-αʹ2 before its dendrites innervate the MB lobes did not influence zonal elaboration of MBON-αʹ3 and MBON-γ2αʹ1 dendrites (**Figs. 2C-F’’ and S5**), indicating that dendritic tiling is not the mechanism by which the borders between MBON dendrites in neighboring zones are determined. Therefore, zonal elaborations of DAN axons and MBON dendrites are largely independent of each other.

**Figure 2.**
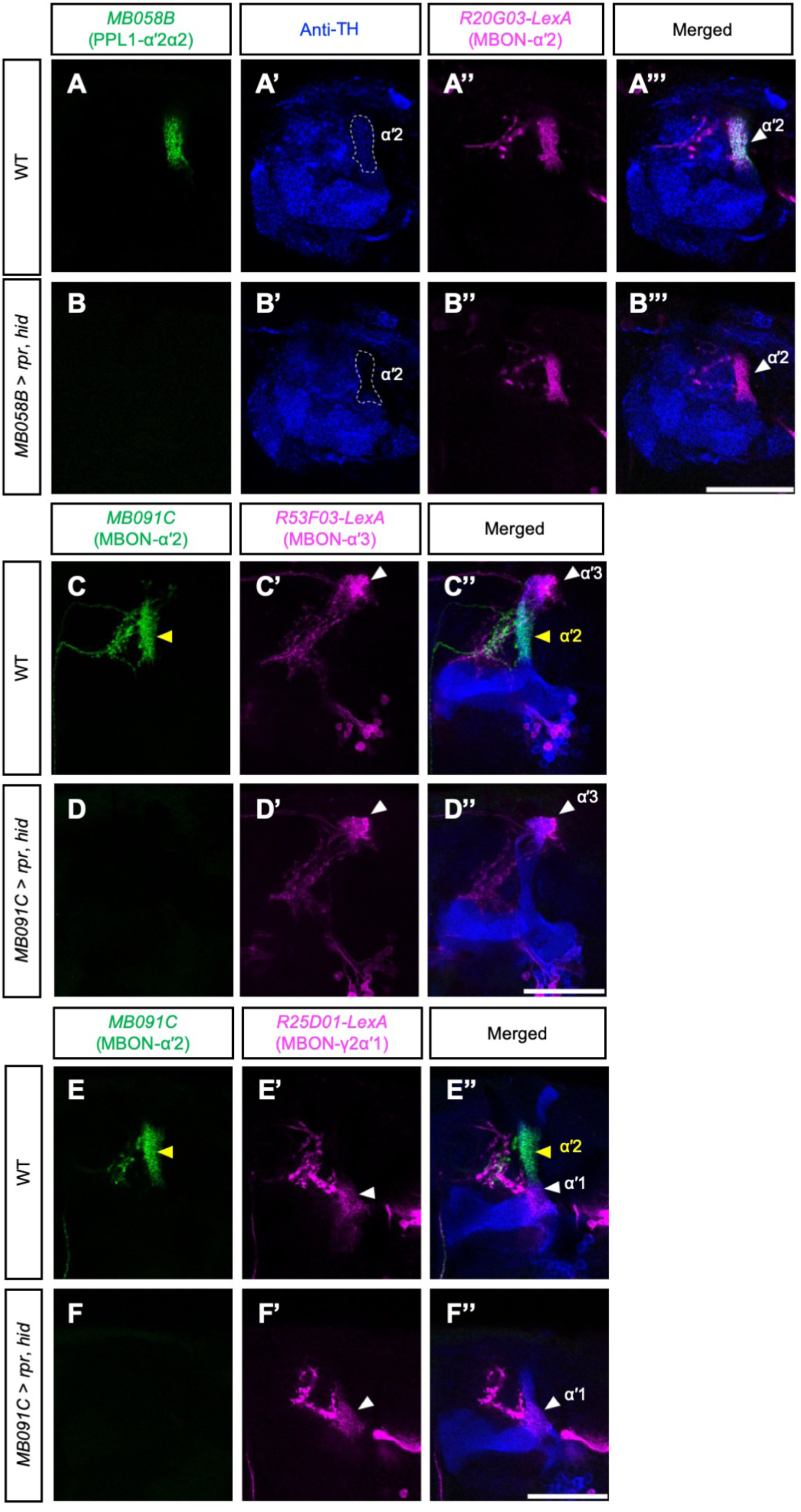
Ablation of PPL1-αʹ2α2 DANs has no effect on MB targeting by neighboring DANs or MBON-αʹ2. **(A-A’’’)** A wild-type brain whose PPL1-αʹ2α2 DANs were labeled with *UAS-mCD8::GFP* driven by *MB058B-splitGAL4* (green, A and A’’’) and whose MBON-αʹ2 was labeled with *LexAop-rCD2::RFP* driven by *R20G03-LexA* (magenta, A’’ and A’’’). The brain was also counterstained with anti-TH antibody (blue, A’ and A’’’). A single focal plane is shown. The white dashed circle in (A’) and the white arrowhead in (A’’’) indicates the αʹ2 zone. **(B-B’’’)** As for (A-A’’’), except that PPL1-αʹ2α2 DANs were ablated with *UAS-rpr,hid* driven by *MB058B-splitGAL4*. **(C-C’’)** A wild-type brain whose MBON-αʹ2 was labeled with *UAS-mCD8::GFP* driven by *MB091C-splitGAL4* (green, C and C’’) and its MBON-αʹ3 labeled with *LexAop-rCD2::RFP* driven by *R53F03-LexA* (magenta, C’ and C’’). The brain was also counterstained with anti-Trio antibody to label the αʹ, βʹ and γ lobes (blue, C’’). The yellow and white arrowheads indicate the αʹ2 and αʹ3 zones, respectively. **(D-D’’)** As for (C-C’’), except that MBON-αʹ2 was ablated using *UAS-rpr,hid* driven by *MB091C-splitGAL4*. **(E-E’’)** A wild-type brain whose MBON-αʹ2 was labeled with *UAS-mCD8::GFP* driven by *MB091C-splitGAL4* (green, E and E’’) and its MBON-γ2αʹ1 labeled with *LexAop-rCD2::RFP* driven by *R25D01-LexA* (magenta, E’ and E’’). The brain was also counterstained with anti-Trio antibody to label the αʹ, βʹ and γ lobes (blue, E’’). A single focal plane is shown. The yellow and white arrowheads indicate the αʹ2 and αʹ1 zones, respectively. **(F-F’’)** As for (E-E’’), except that MBON-αʹ2 was ablated using *UAS-rpr,hid* driven by *MB091C-splitGAL4*. Scale bars: 50 µm. Detailed genotypes are listed in Table S2.

Interestingly, when we ablated MBON-αʹ3, we found that the morphology of the αʹ lobe (labeled by the KC-specific driver *MB247-LexA::p65* and *LexAop-rCD2::RFP*) was altered (**Fig. S3**) in that it became shorter and the tip of the αʹ lobe (i.e. the αʹ3 zone) became smaller in adult brains lacking MBON-αʹ3 (**Fig. S3A-E**). We followed the development of the MB lobes at different pupal stages in MBON-αʹ3-ablated brains and found that growth of the αʹ lobe was slower than for wild-type controls (**Figs. S7 and S3F**). This result suggests that the shorter αʹ3 tip was caused by a developmental defect, not neurodegeneration. The αʹ3 zone did not completely disappear when MBON-αʹ3 was ablated because when we labeled MBON-αʹ2 with *R20G03-LexA::P65*, its dendrites remained clear of the reduced tip of the αʹ lobe (**Fig. S3G-H’’**). These data suggest that the MBON-αʹ3 dendrites might provide a positive signal to promote growth of the KC axons. Alternatively, innervation by the MBON-αʹ3 dendrites might simply inflate the αʹ3 zone in wild-type brains. However, this latter explanation seems less likely because we did not observe a similar reduction of the αʹ2 zone when MBON-αʹ2 dendrites were ablated (**Fig. S3I**).

### KC axons are essential for zonal elaborations of DAN axons

To investigate the role of KC axons in axonal and dendritic targeting of DANs and MBONs, respectively, we took advantage of the *alpha lobe absent* (*ala*) fly mutant. Brains of the *ala* mutant have been shown to randomly lose vertical or horizontal lobes of the MB (Pascual and Préat, 2001; Pascual et al., 2005). We found that in *ala* mutants that lose MB vertical lobes on one side of the brain, both contralateral and ipsilateral PPL1-αʹ2α2 axons wandered around the areas near their missing targets without forming normal zonal elaboration patterns (**Fig. 3A-B’’**). Mistargeting of PPL1-αʹ2α2 axons was completely correlated with the lobe-missing phenotype (n=9), so it was not caused by loss of the *ala* gene *per se* but was a consequence of the missing lobe. We also examined PPL1-αʹ2α2 axons in the *ala* mutant brains at 30 h APF, when obvious zonal innervation by the axons is first observed (**Fig. 1C-C’**). We found that when vertical lobes were absent, PPL1-αʹ2α2 axons failed to form a zonal network from the very beginning of the elaboration process, and some mistargeted neurites could be found slightly above and posterior to where the αʹ2 and α2 zones are supposed to be located (**Fig. 3C-D’**). Since these mistargeted axons could still project to sites close to their normal targets (**Fig. 3**), KC axons do not appear to provide long-range guidance cues for these axons. Instead, the KC axons may provide local positional cues for the DAN axons to form zonal networks. This latter scenario was further evidenced when we used a multi-color flip-out technique to label PPL1-αʹ2α2 and PPL1-αʹ3 DANs in different colors. We found that the primary neurites of these two DANs entered the MB lobes at a similar location (**Fig. S8**). However, PPL1-αʹ2α2 axons exclusively occupied the αʹ2 and α2 zones below the entry point, whereas PPL1-αʹ3 axons specifically covered the αʹ3 zone above the entry point (**Fig. S8**), suggesting local cues exist that direct zonal elaboration of DAN axons.

**Figure 3.**
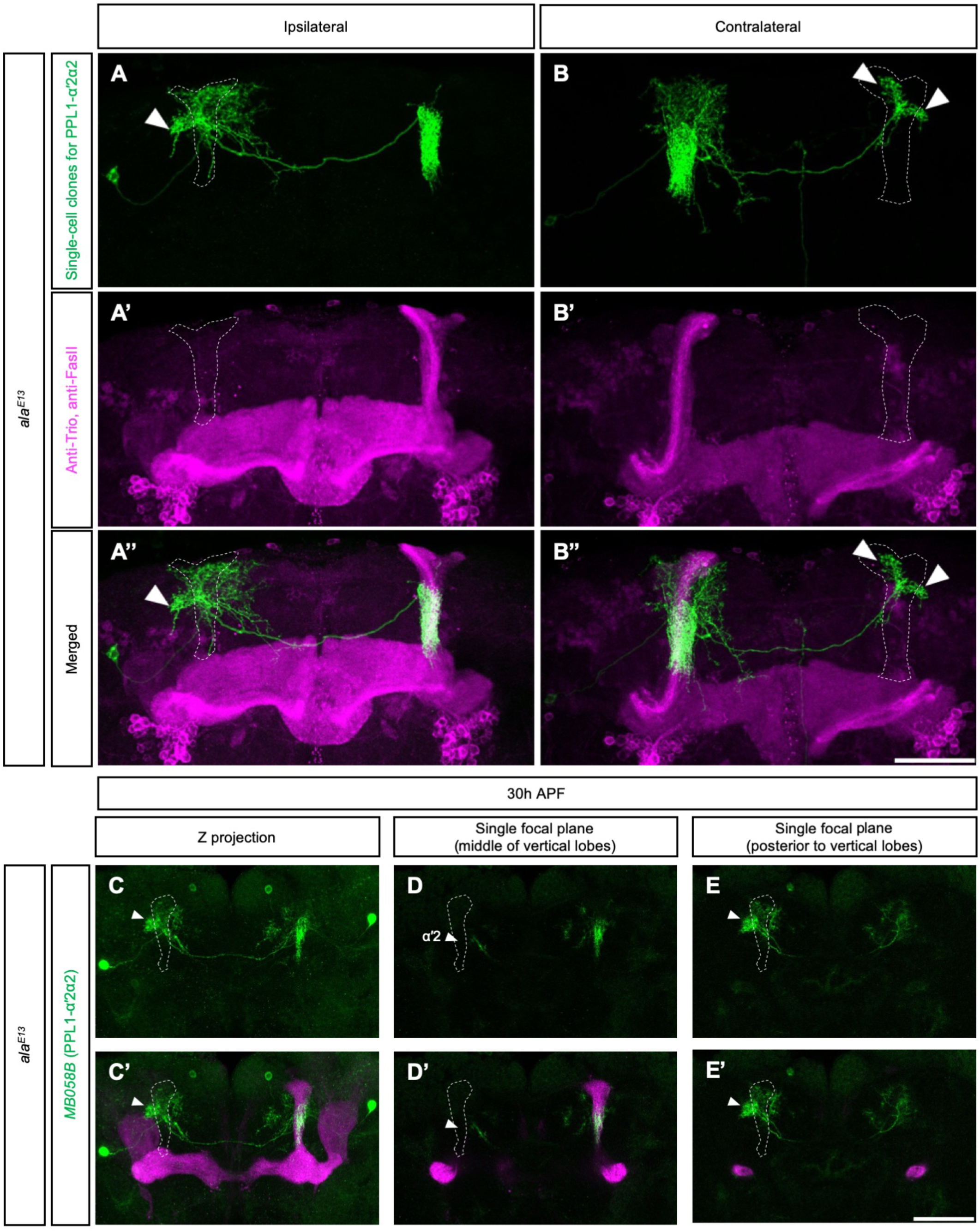
Loss of vertical lobes results in mistargeted PPL1-αʹ2α2 axons. **(A-B’’)** Single-cell flip-out clones of PPL1-αʹ2α2 DANs (green; A, A’’, B, and B’’) in *ala* mutant brains from which the MB has lost its vertical lobes on the ipsilateral (A-A’’) or contralateral (B-B’’) side to the clones. The brains were counterstained with anti-Trio and anti-FasII antibodies to label all MB lobes (magenta; A’-A’’ and B’-B’’). The white arrowheads in (A, A’’, B and B’’) indicate the mistargeted axons. **(C-C’)** Axonal innervation patterns of PPL1-αʹ2α2 DANs in an *ala* mutant brain at 30 h APF. The PPL1-αʹ2α2 DANs were labeled with *UAS-mCD8::GFP* driven by *MB058B-splitGAL4* (green), and all KCs were labeled with *LexAop-rCD2::RFP* driven by *MB247-LexA::p65* (magenta, C’). The vertical lobes were missing in one hemibrain. The white arrowheads indicate the mistargeted axons near the missing lobes. **(D-D’)** A single focal plane from (C-C’) showing the αʹ2 zones. The white arrowheads indicate the position of the αʹ2 zone. **(E-E’)** A single focal plane from (C-C’) showing the layer immediately posterior to the vertical lobes. The white arrowheads indicate the mistargeted axons locate laterally and dorsally to the putative missing αʹ2/α2 zones. White dashed lines outline the putative position of the missing lobes. Scale bars: 50 µm. Detailed genotypes are listed in Table S2.

### Competition between MB lobe zones for MBON-αʹ2 dendrites

Surprisingly, when the MB vertical lobes were missing, MBON-αʹ2 dendrites targeted the βʹ2mp and αʹ1 zones instead (**Fig. 4A-B’’**). A previous study showed that in *ala* mutants lacking vertical MB lobes, approximately half of the KCs lose their vertical branches and the vertical branches of the remaining half project horizontally (Pascual et al., 2005). However, the mistargeting of MBON-αʹ2 we report here is unlikely caused by the horizontally-projecting KC vertical branches for two reasons. First, the zones innervated by the mistargeted MBON-αʹ2 dendrites are located at either end, but not the middle, of the βʹ lobe. Second, the guidance molecules required for MBON-αʹ2 to innervate the αʹ2 zone are different from those required for it to innervate the βʹ2 and αʹ1 zones (see below). When we followed the development of the MBON-αʹ2 dendrites in the absence of their normal target, we found that they never formed a zonal network near the missing target, but started to innervate the βʹ2mp and αʹ1 zones at around 72 h APF (**Fig. 4C-E’**). Taken together, these results indicate that there are competing guidance cues from multiple MB lobe zones for the MBON-αʹ2 dendrites. When the stronger guidance cues from the αʹ2 zones are missing, the MBON-αʹ2 dendrites are attracted to other zones.

**Figure 4.**
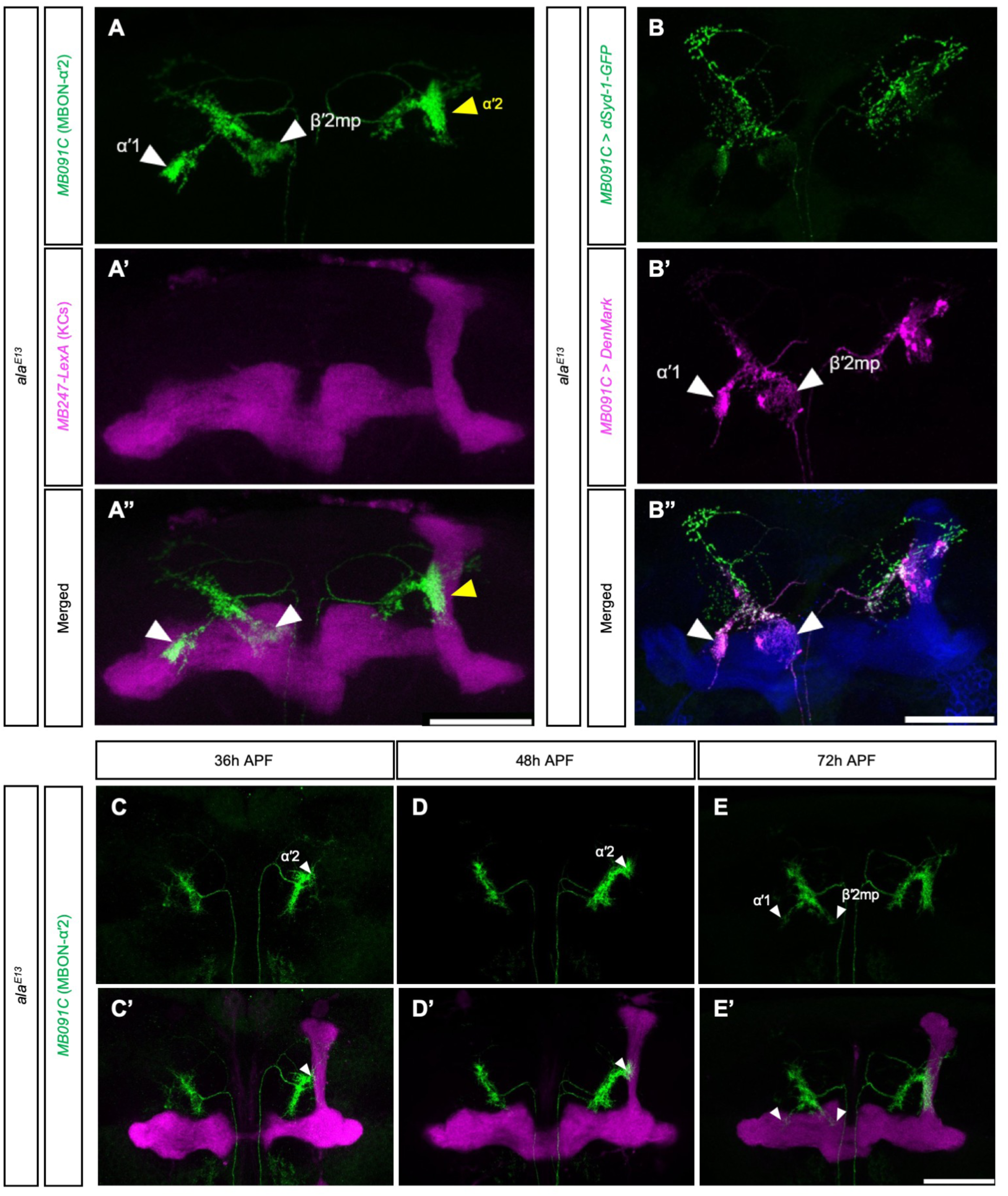
MBON-αʹ2 dendrites innervate other MB lobe zones when their normal target zone is missing. **(A-A’’)** Innervation pattern of MBON-αʹ2 dendrites labeled with *UAS-mCD8::GFP* driven by *MB091C-splitGAL4* (green, A and A’’) in an *ala* mutant brain from which the vertical lobes are missing in one hemibrain. The KCs were labeled with *LexAop-rCD2::RFP* driven by *MB247-LexA::p65* (magenta, A’ and A’’). White arrowheads in (A and A’’) indicate the αʹ1 and βʹ2mp zones. Yellow arrowheads in (A and A’’) indicate the normal αʹ2 zone. **(B-B’’)** As for (A-A’’), except that the MBON-αʹ2 was labeled with *UAS-dSyd-1-GFP* (green, presynaptic marker, B and B’’) and *UAS-DenMark* (magenta, postsynaptic marker, B’ and B’’). White arrowheads in (B’ and B’’) indicate the αʹ1 and βʹ2mp zones. **(C-E’)** Targeting of MBON-αʹ2 dendrites labeled with *UAS-mCD8::GFP* driven by *MB091C-splitGAL4* (green) in *ala* mutant brains at 36 (C-C’), 48 (D-D’), and 72 (E-E’) h APF. The KCs were labeled with *LexAop-rCD2::RFP* driven by *MB247-LexA::p65* (magenta, C’, D’ and E’). Vertical lobes were missing from one hemibrain. The white arrowheads in (C-D’) indicate the αʹ2 zone, and the arrowheads in (E-E’) indicate ectopic innervations of the MBON-αʹ2 dendrites. Scale bars: 50 µm. Detailed genotypes are listed in Table S2.

### Overexpression of *sema1a* in DANs mistargets their dendrites to specific MB lobe zones

During the course of identifying genes that control MB circuit assembly, we found that when *sema1a* was overexpressed in PPL1-αʹ2α2 DANs, their dendrites that normally elaborate loosely behind the MB vertical lobes (**Fig. 5A-A’’**) instead innervated the βʹ2p, αʹ1, and αʹ3 zones (**Fig. 5B-B’’**). We confirmed that these misguided neurites are dendrites by labeling PPL1-αʹ2α2 with DenMark and dSyd1-GFP (**Fig. 5E-E’’**). The dSyd-1-GFP-labeled axons were not affected by overexpression of *sema1a*. Sema1a can function as a ligand or receptor in different developmental contexts (Cafferty et al., 2006; Godenschwege et al., 2002; Komiyama et al., 2007; Winberg et al., 1998). We found that overexpression in PPL1-αʹ2α2 DANs of *sema1a* lacking its intracellular domain (Sema1a-mEC-5xmyc and Sema1a-mEC.Fc-5xmyc) (Jeong et al., 2012) had no effect on their dendrites (**Fig. 5C-D’’**), suggesting that when overexpressed, Sema1a works as a receptor to guide PPL1-αʹ2α2 dendrites to the MB lobe zones where the ligands for Sema1a are likely produced. Overexpression of *sema1a* in PPL1-γ2αʹ1 DANs using *MB296B-splitGAL4* also caused mistargeted dendrites, but in this instance they innervated the βʹ2mp and βʹ2a zones (**Fig.5F-I’’**). It is striking that overexpression of *sema1a* in different DANs targets their dendrites to distinct βʹ2 zones, suggesting that Sema1a may work with other guidance molecules to determine zonal specificity for different dendrites.

**Figure 5.**
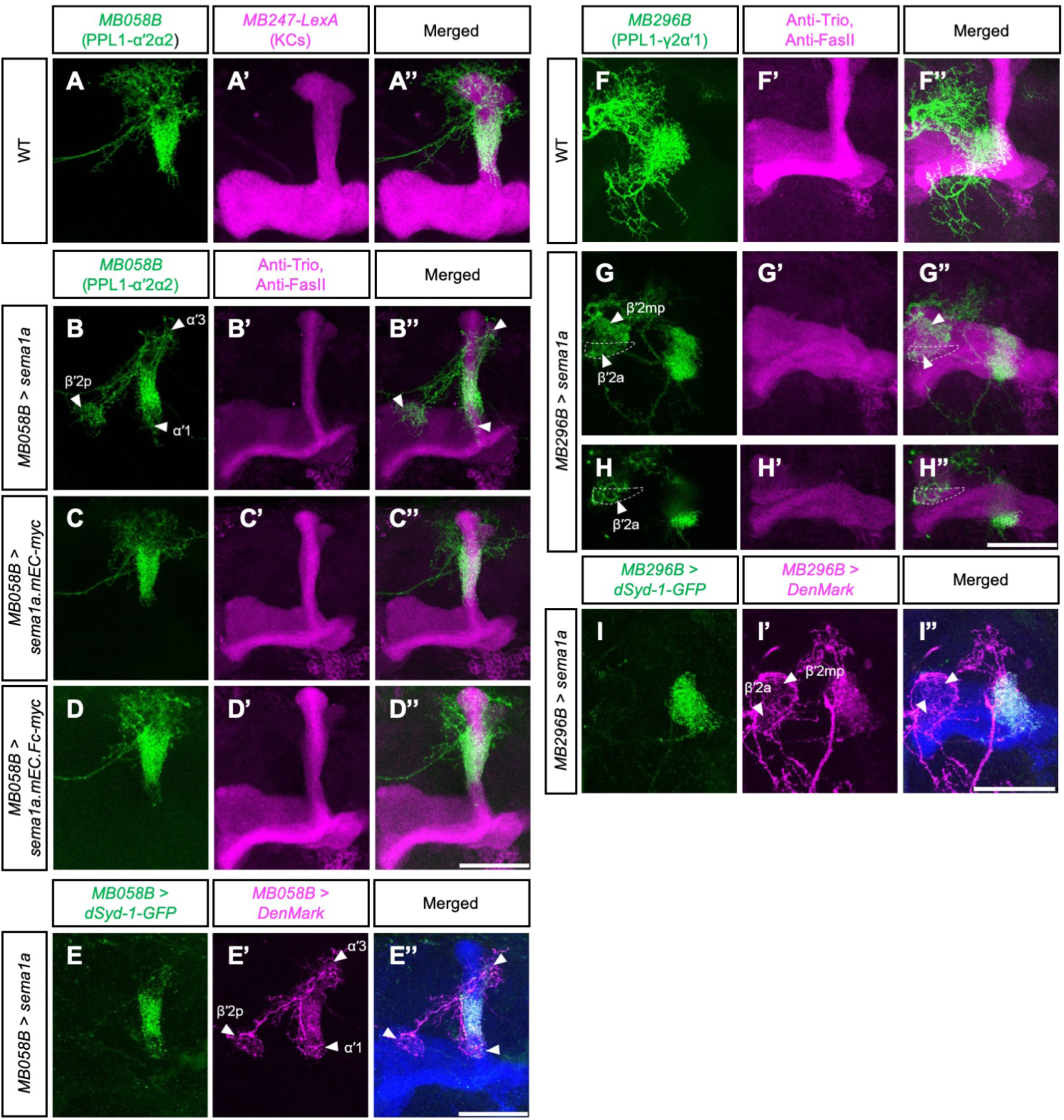
Overexpression of *sema1a* redirects PPL1-αʹ2α2 and PPL1-γ2αʹ1 dendrites to distinct MB lobe zones. **(A-A’’)** PPL1-αʹ2α2 DANs labeled with *UAS-mCD8::GFP* driven by *MB058B-splitGAL4* (green, A and A’’) in a wild-type brain. The brain was counterstained with anti-Trio and anti-FasII antibodies to label all MB lobes (magenta, A’ and A’’). **(B-B’’)** As for (A-A’’), except that Sema1a was additionally overexpressed in PPL1-αʹ2α2 DANs using *UAS-sema1a* driven by *MB058B-splitGAL4*. Arrowheads in (B and B’’) indicate the βʹ2p, αʹ1, and αʹ3 zones. **(C-C’’)** As for (B-B’’), but *UAS-sema1a* was replaced with *UAS-sema1.mEC-5xmyc*. **(D-D’’)** As for (B-B’’), but *UAS-sema1a* was replaced with *UAS-sema1.mEC.Fc-5xmyc*. **(E-E’’)** As for (B-B’’), except that the PPL1-αʹ2α2 DANs were labeled with *UAS-dSyd-1-GFP* (green, presynaptic marker, E and E’’) and *UAS-DenMark* (magenta, postsynaptic marker, E’ and E’’). Arrowheads in (E’ and E’’) indicate the βʹ2p, αʹ1, and αʹ3 zones. **(F-F’’)** PPL1-γ2αʹ1 DANs labeled with *UAS-mCD8::GFP* driven by *MB296B-splitGAL4* (green, F and F’’) in a wild-type brain. The brain was counterstained with anti-Trio and anti-FasII antibodies to label all MB lobes (magenta, F’ and F’’). **(G-G’’)** As for (F-F’’), except that Sema1a was additionally overexpressed in PPL1-γ2αʹ1 DANs using *UAS-sema1a* driven by *MB296B-splitGAL4*. Arrowheads in (G and G’’) indicate the βʹ2mp and βʹ2a zones. Note that this manipulation resulted in loss of MB vertical lobes due to an unknown mechanism. **(H-H’’)** A single focal plane from (G-G’’) showing the innervation of ectopic PPL1-γ2αʹ1 neurites in the βʹ2a zone (outlined by white dashed lines). **(I-I’’)** As for (G-G’’), except that the PPL1-γ2αʹ1 DANs were labeled with *UAS-dSyd-1-GFP* (green, I and I’’) and *UAS-DenMark* (magenta, I’ and I’’). Arrowheads in (I’ and I’’) indicate the βʹ2mp and βʹ2a zones. Scale bars: 50 µm. Detailed genotypes are listed in Table S2.

Overexpression of *sema1a* cannot always outcompete the endogenous dendritic targeting program, as is evident for MBON-αʹ2 in which overexpression of *sema1a* did not lead to any dendritic targeting phenotype (**Fig. S9**). However, ectopic innervation by *sema1a*-overexpressed PPL1-αʹ2α2 and PPL1-γ2αʹ1 dendrites in the βʹ2mp and αʹ1 zones is reminiscent of the mistargeting of MBON-αʹ2 dendrites in the absence of MB vertical lobes (**Fig. 4**). Therefore, we tested if *sema1a* is required for the mistargeting of MBON-αʹ2 dendrites in *ala* mutant brains. We knocked down *sema1a* from MBON-αʹ2 in *ala* mutant flies and found that the MBON-αʹ2 dendrites with reduced *sema1a* expression no longer innervated the βʹ2mp and αʹ1 zones when their normal target was missing (**Fig. 6**). However, knockdown of *sema1a* did not affect MBON-αʹ2 dendritic targeting to the αʹ2 zone in the normal vertical lobes (**Fig. 6**). These results suggest that MBON-αʹ2 may express *sema1a* as well as a yet unidentified guidance molecule. In wild-type brains, this unknown guidance molecule outcompetes *sema1a* to target MBON-αʹ2 dendrites to the αʹ2 zone. However, when the αʹ2 zone is missing, a *sema1a*-depedent guidance mechanism takes precedence, directing the MBON-αʹ2 dendrites to the βʹ2mp and αʹ1 zones.

**Figure 6.**
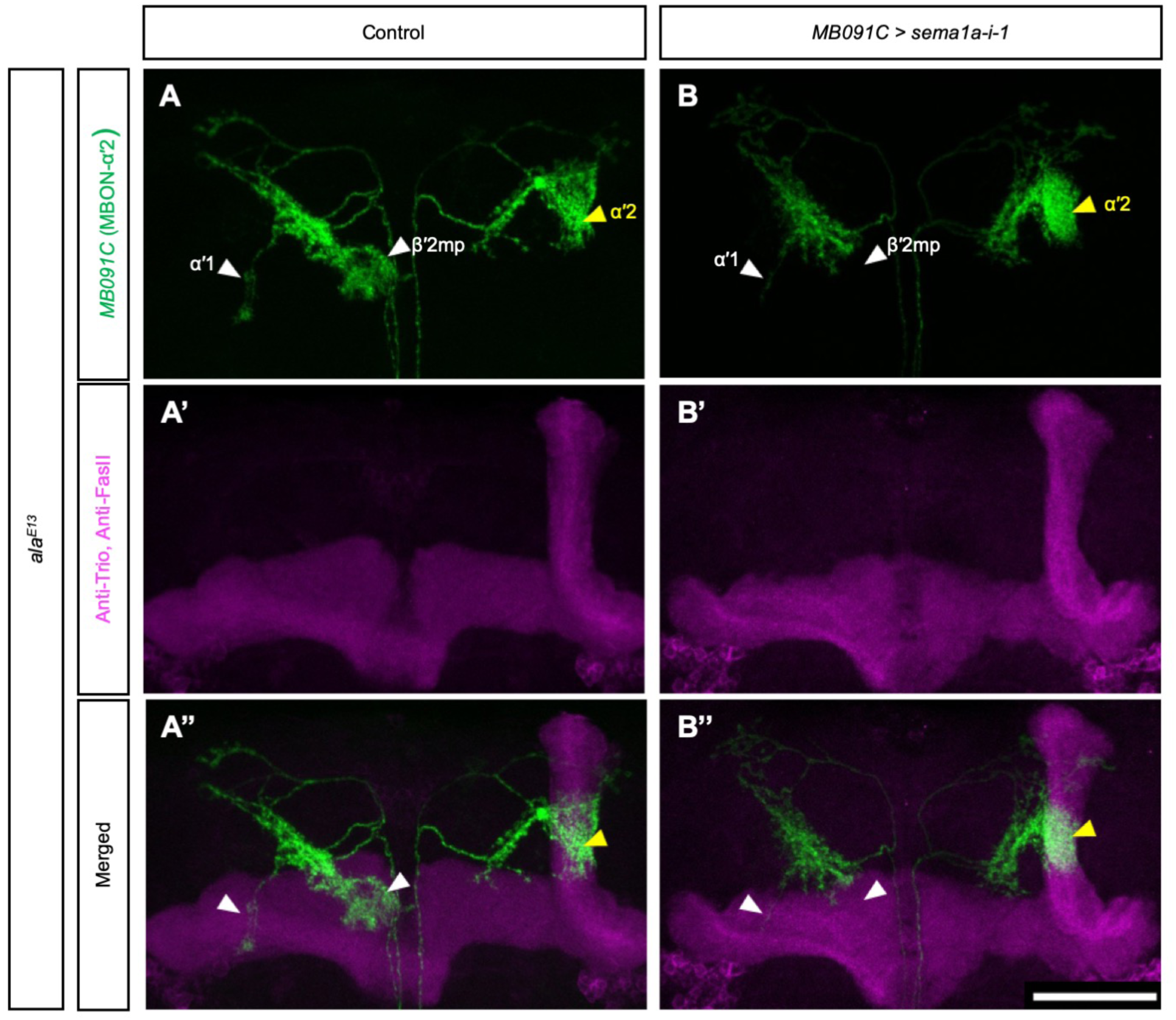
*sema1a* is required for the mistargeting of MBON-αʹ2 dendrites due to missing vertical lobes. **(A-A’’)** The innervation pattern of MBON-αʹ2 dendrites labeled with *UAS-mCD8::GFP* driven by *MB091C-splitGAL4* (green, A and A’’) in an *ala* mutant brain lacking MB vertical lobes in one hemibrain. All MB lobes were labeled with *LexAop-rCD2::RFP* driven by *MB247-LexA::p65* (magenta, A’ and A’’). **(B-B’’)** As for (A-A’’), except that *UAS-sema1a-RNAi* was additionally expressed in MBON-αʹ2. The white arrowheads indicate the mistargeted dendrites in the βʹ2mp and αʹ1 zones and the yellow arrowheads indicate the normal innervation in the αʹ2 zones. Note the greatly reduced ectopic dendritic innervations and that normal αʹ2 innervation is not affected by *sema1a* knockdown. Scale bars: 50 µm. Detailed genotypes are listed in Table S2.

### *Sema1a* is required for dendritic targeting of several MBONs

The ectopic innervation of *sema1a*-overexpressed PPL1-αʹ2α2 and PPL1-γ2αʹ1 dendrites in the βʹ2, αʹ1, and αʹ3 zones prompted us to examine the role of *sema1a* in dendritic targeting of the MBONs innervating these three zones. We found that RNAi knockdown of *sema1a* in MBON-β2βʹ2a using *MB399C-splitGAL4* resulted in mistargeting of the dendrites that normally innervate the βʹ2a zone (**Fig. 7A-D**). Instead of innervating the βʹ2a zone, these dendrites projected upwards to regions near the central complex. The MBON-β2βʹ2a dendrites innervating the β2 zone were minimally affected by *sema1a* knockdown (**Fig. 7A-D**), suggesting that *sema1a* regulates dendritic targeting of MBON-β2βʹ2a in a zone-specific manner. Similarly, when *sema1a* was knocked down in MBON-αʹ3 using *MB027B-splitGAL4*, the MBON-αʹ3 dendrites failed to concentrate in the αʹ3 zone but elaborated loosely around the αʹ3 zone (**Fig. 7E-H**). We confirmed these findings by creating *sema1a* MARCM (mosaic analysis with a repressible cell marker; (Lee and Luo, 1999)) clones for MBON-β2βʹ2a (**Fig. 8A-B’**) and MBON-αʹ3 (**Fig. 8C-D’**). The dendritic targeting defects observed in these *sema1a*-mutant MBON-β2βʹ2a and MBON-αʹ3 were consistent with the phenotypes caused by *sema1a* RNAi knockdown. To confirm that MBON-β2βʹ2a and MBON-αʹ3 express *sema1a*, we used *sema1a* Artificial exon that produces V5-tagged Sema1a and LexA::p65 under the control of the endogenous *sema1a* enhancer when an FRT-stop-FRT cassette in the exon is removed by Flipase (FLP) (Pecot et al., 2013). Consistently, V5-tagged Sema1a and LexA::P65 driven by the *sema1a* enhancer were detected in MBON-β2βʹ2a and MBON-αʹ3 when *MB399C-splitGAL4* and *MB027B-splitGAL4* were used, respectively, to drive *UAS-FLP* (**Fig. S10**).

**Figure 7.**
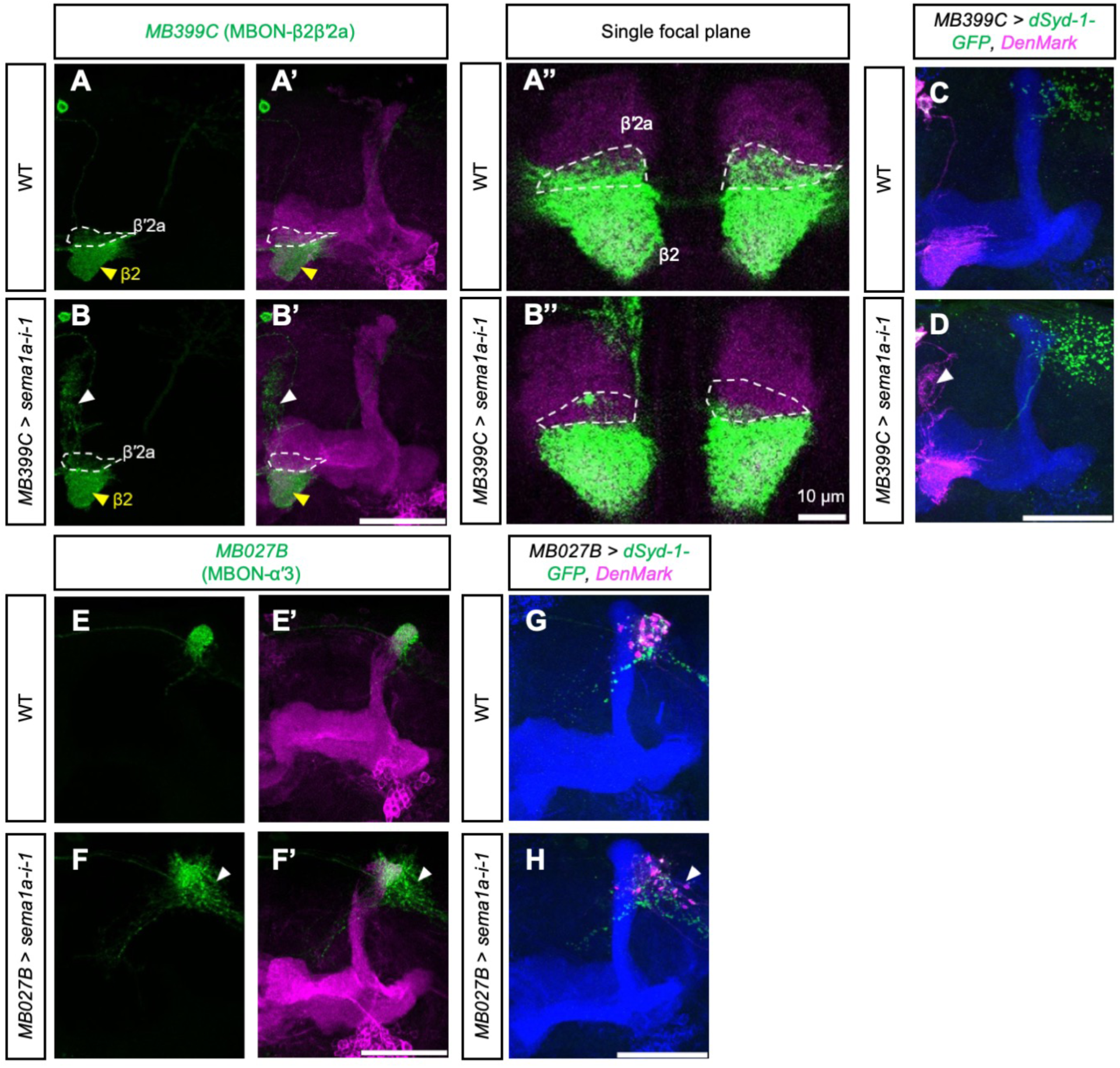
*sema1a* is required for the dendritic targeting of MBON-β2βʹ2a and MBON-αʹ3. **(A-A’’)** The innervation pattern of MBON-β2βʹ2a dendrites labeled with *UAS-mCD8::GFP* driven by *MB399C-splitGAL4* (green, A-A’’). The αʹ, βʹ and γ lobes were labeled with anti-Trio antibody (magenta, A’ and A’’). The βʹ2a zones are outlined with white dashed circles, and the β2 zones are indicated by yellow arrowheads. (A’’) is a high magnification view of a single focal plane showing the βʹ2a and β2 zones. **(B-B’’)** As for (A-A’’), except that *UAS-sema1a-RNAi-1* was additionally expressed by *MB399C-splitGAL4* in MBON-β2βʹ2a. The white arrowheads in (B and B’) indicate the mistargeted neurites. **(C-D)** Wild-type (C) and *UAS-sema1a-RNAi-1*-expressing (D) MBON-β2βʹ2a labeled with dSyd-1-GFP (green, presynaptic marker) and DenMark (magenta, postsynaptic marker) driven by *MB399C-splitGAL4*. The brains were counterstained with anti-Trio and anti-FasII antibodies to label all MB lobes (blue). The white arrowhead in (D) indicates the mistargeted dendrites. **(E-F’)** Wild-type (E-E’) and *UAS-sema1a-RNAi*-1-expressing (F-F’) MBON-αʹ3 labeled with *UAS-mCD8::GFP* driven by *MB027B-splitGAL4* (green). The brains were counterstained with anti-Trio antibody to label the αʹ, βʹ and γ lobes (magenta, E’ and F’). The white arrowheads in (F and F’) indicate the mistargeted dendrites. **(G-H)** Wild-type (G) and *UAS-sema1a-RNAi-1*-expressing (H) MBON-αʹ3 labeled with dSyd-1-GFP (green, presynaptic marker) and DenMark (magenta, postsynaptic marker) driven by *MB027B-splitGAL4*. The brains were counterstained with anti-Trio and anti-FasII antibodies to label all MB lobes (blue). The white arrowhead in (H) indicates the mistargeted dendrites.

**Figure 8.**
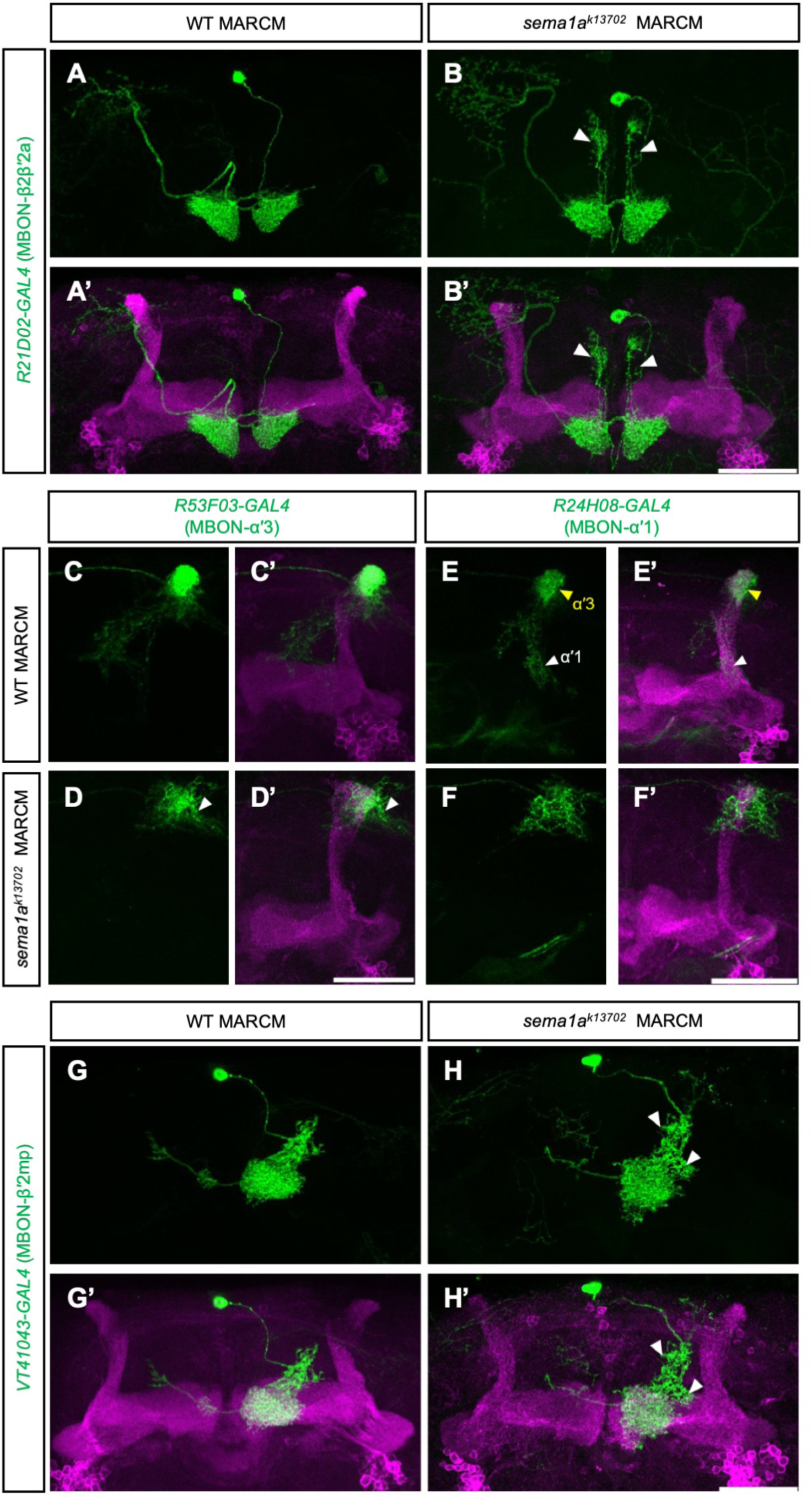
*sema1a* mutant MARCM clones for MBON-β2βʹ2a, MBON-αʹ3, MBON-αʹ1, and MBON-βʹ2mp exhibit defective dendritic targeting. **(A-B’)** Wild-type (A-A’) and *sema1a^k13702^* (B-B’) MARCM clones for MBON-β2βʹ2a labeled with *UAS-mCD8::GFP* driven by *R21D02-GAL4* (green). The brains were counterstained with anti-Trio antibody to label the αʹ, βʹ and γ lobes (magenta, A’ and B’). The white arrowheads in (B-B’) indicate the mistargeted dendrites. **(C-D’)** Wild-type (C-C’) and *sema1a^k13702^* (D-D’) MARCM clones for MBON-αʹ3 labeled with *UAS-mCD8::GFP* driven by *R53F03-GAL4* (green). The brains were counterstained with anti-Trio antibody to label the αʹ, βʹ and γ lobes (magenta, C’ and D’). The white arrowhead in (D-D’) indicates the mistargeted dendrites. **(E-F’)** Wild-type (E-E’) and *sema1a^k13702^* (F-F’) MARCM clones for MBON-αʹ3 and MBON-αʹ1 labeled with *UAS-mCD8::GFP* driven by *R24H08-GAL4* (green). The brains were counterstained with anti-Trio antibody to label the αʹ, βʹ and γ lobes (magenta, E’ and F’). The yellow arrowheads in (E-E’) indicate the innervation of MBON-αʹ3 and the white arrowheads in the same images indicate the innervation of MBON-αʹ1. Note that MBON-αʹ3 was always accompanied by MBON-αʹ1 in the wild-type MARCM clones (n = 17), indicating that they are from the same lineage. MBON-αʹ3 and MBON-αʹ1 dendritic innervations were both defective in *sema1a^k13702^* MARCM clones (F-F’). **(G-H’)** Wild-type (G-G’) and *sema1a^k13702^* (H-H’) MARCM clones for MBON-βʹ2mp labeled with *UAS-mCD8::GFP* driven by *VT41043-GAL4* (green). The brains were counterstained with anti-Trio antibody to label the αʹ, βʹ and γ lobes (magenta, G’ and H’). The white arrowheads in (H-H’) indicate the mistargeted dendrites. Scale bars: 50 µm. Detailed genotypes are listed in Table S2.

MARCM also allowed us to test the role of *sema1a* in MBONs for which we do not have GAL4 lines to drive RNAi sufficiently early during their development. Apart from innervation by MBON-β2βʹ2a dendrites, the βʹ2 zone is also innervated by MBON-βʹ2mp and MBON-γ5βʹ2a dendrites, and the αʹ1 zone is further innervated by MBON-γ2αʹ1 and MBON-αʹ1 dendrites. We found that dendritic targeting of MBON-γ2αʹ1 and MBON-γ5βʹ2a remained normal in *sema1a* MARCM clones (**Fig. S11**). However, *sema1a* MARCM clones for MBON-αʹ1 exhibited a severe dendritic targeting defect (**Fig. 8E-F’**). We induced clones at 4-8 h after egg laying (AEL) and labeled the MBON-αʹ1 dendrites using *R24H08-GAL4*, which also labels MBON-αʹ3 dendrites. Interestingly, all of the MBON-αʹ1 clones were accompanied by MBON-αʹ3 clones, suggesting that these two neuron types are from the same neuronal lineage. We also observed a dendritic targeting defect in *sema1a* MARCM clones for MBON-βʹ2mp (**Fig. 8G-H’**). Taken together, these results indicate that *sema1a* is required for dendritic targeting in most MBONs that innervate the βʹ2, αʹ1, and αʹ3 zones.

### Canonical partners of Sema1a are not essential for the targeting of MBON dendrites

Sema2a, Sema2b, and PlexA have been identified as ligands for Sema1a (Sweeney et al., 2011; Yu et al., 2010). To test if these molecules play a role in the targeting of MBON dendrites, we first examined the dendritic innervation patterns of MBON-β2βʹ2a and MBON-αʹ3 in flies carrying homozygous mutations in both *sema2a* and *sema2b* genes (*Sema2b^f02042^, sema2a^p2^*) (Sweeney et al., 2011). Although MB lobe structure is slightly abnormal in these flies, morphology of MBON-β2βʹ2a and MBON-αʹ3 dendrites and their zonal innervations were essentially normal (**Fig. S12A-B’**). Therefore, Sema2a and Sema2b are unlikely to be the guidance molecules that direct *sema1a*-positive MBON dendrites to specific MB lobe zones. Flies lacking PlexA activity fail to survive to adulthood. To examine the potential role of PlexA in guiding the MBON dendrites, we took advantage of a weaker *sema1a-RNAi-2* line whose overexpression resulted in mild targeting defects of MBON-αʹ3 and MBON-β2βʹ2a dendrites (**Fig. S12C-C’ and E-E’**) and then checked if loss of one *plexA* allele exacerbated the defects. However, the targeting defects of MBON-β2βʹ2a and MBON-αʹ3 dendrites were not exacerbated in flies heterozygous for a *plexA* mutant allele, *plexA^EY16548^* (**Fig. S12C-F’**). We also expressed an *UAS-plexA-RNAi* construct that has been shown to knock down *plexA* effectively in all neurons (*nSyb-GAL4*) or in all glia (*repo-GAL4*) (Pecot et al., 2013). We then examined the innervation by MBON-β2βʹ2a and MBON-βʹ2mp (*R12C11-LexA*-driven rCD2-RFP) and MBON-αʹ3 (*R53F03-LexA*-driven rCD2-RFP) dendrites in the MB lobes. However, we did not observe any dendritic targeting phenotype in all cases (**Fig. S12G-N’**). Therefore, there is no evidence that the three known Sema1a ligands are involved in guiding MBON dendrites during assembly of the MB circuit.

## Discussion

The MB is the most studied structure in the fly brain. It is conventionally considered as an olfactory learning and memory center (Aso et al., 2014; de Belle and Heisenberg, 1994; Heisenberg, 2003). Recently, it has also been shown to receive other sensory inputs and to mediate motivational control of innate behavior (Krashes et al., 2009; Masek and Scott, 2010; Masek et al., 2015; Senapati et al., 2019; Tsao et al., 2018; Vogt et al., 2014, 2016). Almost all the neurons that form the MB circuit have been identified, and genetic means of labeling individual neuronal types have been developed (Aso et al., 2014). However, our understanding of the assembly process of this important structure is essentially lacking. In this study, we provide an initial characterization of this process and reveal several interesting principles.

The most intriguing feature of the organization of the MB circuit is the zonal innervation of the MB lobes by DAN axons and MBON dendrites. Electron microscopy reconstruction of the MB lobes shows that in each zone the DAN axons and MBON dendrites form synapses with each other and the KC axons (Takemura et al., 2017). The borders of the zones are clear, and there is minimal overlap between DAN axons or MBON dendrites in the neighboring zones. With such a highly organized neural network, elaborate interactions among the extrinsic neurons might be expected. For example, dendritic tiling, as observed between dendritic arborization (da) neurons in fly larvae, might be required for the formation of zonal borders (Grueber et al., 2002), and matching between DAN axons and MBON dendrites in the same zone might be important for these neurites to establish proper zonal innervation patterns. However, our results suggest that the targeting and elaboration networks of DAN axons and MBON dendrites are largely independent. In the αʹ2 zone, where innervation by DAN axons precedes that by MBON dendrites, ablation of DANs does not affect zonal elaboration of the MBON dendrites. Moreover, upon ablation of one type of DAN or MBON, morphologies of the neighboring neurites appear to be normal. Therefore, the extent and location of zonal network elaboration by DAN axons and MBON dendrites might be primarily determined by the interactions between KC axons and these extrinsic neurons.

The KC axons are apparently essential for zonal arborization of the DAN axons and MBON dendrites. When their target lobes are absent, DAN axons wander around the missing target and generate less extensive neurite arbors. Therefore, the KC axons appear to promote growth and arborization of DAN axons. Do the KC axons also determine the extent and borders of the zonal arbors? It is still possible that the KC axons simply provide an anchoring point on which DAN axons and MBON dendrites grow, and that the extent of their arborization is determined cell-autonomously as an intrinsic property. However, the axonal innervation pattern of PPL1 DANs argues against this latter possibility. The axonal projections of PPL1-αʹ3 and PPL1-αʹ2α2 DANs enter the MB vertical lobes at almost the same location, but specifically innervate distinct zones on opposite sides of the entry point. Therefore, different segments in the KC axons might provide positional cues for the DANs and MBONs, although direct evidence for this possibility awaits investigation. If they exist, what could be the nature of these positional cues? One possibility is that the KC axonal segment in each zone expresses a unique guidance molecule that binds to a receptor on the neurites of the DANs and MBONs projecting to that segment. However, when the target zone of MBON-αʹ2 is lost, its dendrites instead innervate the βʹ2mp and αʹ1 zones. This result suggests that an MBON can respond to multiple positional cues. Furthermore, since *sema1a* is required for MBONs to innervate multiple specific MB lobe zones, the positional cues are likely combinatorial.

Sema1a can function as both a ligand and a receptor (Cafferty et al., 2006; Godenschwege et al., 2002; Komiyama et al., 2007; Winberg et al., 1998). Using *sema1a* transgenes that lack the intracellular domain, we demonstrated that it works as a receptor in MBONs to guide their dendrites. PlexA and Sema2a/2b are known ligands for Sema1a (Sweeney et al., 2011; Yu et al., 2010). However, dendritic targeting by MBONs was minimally affected in flies homozygous for *sema2a/2b* double mutants or when we knocked down PlexA either pan-neuronally or in glia. Moreover, loss of one copy of *plexA* did not exacerbate the dendritic targeting defect caused by RNAi knockdown of *sema1a* in MBONs. Therefore, these canonical Sema1a ligands do not seem to play a role in guiding MBON dendrite targeting. Thus, the molecular mechanisms that direct *sema1a*-positive MBON dendrites to specific MB lobe zones remain unknown.

## Experimental procedures

### Fly strains and husbandry

Fly lines used in this study are: (1) *MB077C-splitGAL4* (Bloomington Drosophila Stock Center, BDSC:68284) (Aso et al., 2014); (2) *MB091C-splitGAL4* (gift from Y. Aso) (Aso et al., 2014). *MB027B-splitGAL4* (BDSC:68301) (Aso et al., 2014); (3) *MB058B-splitGAL4* (BDSC:68278) (Aso et al., 2014); (4) *MB296B-splitGAL4* (BDSC:68308) (Aso et al., 2014); (5) *MB399C-splitGAL4* (gift from Y. Aso) (Aso et al., 2014); (6) *R53F03-GAL4* (BDSC:48197) (Jenett et al., 2012); (7) *R21D02-GAL4* (BDSC:48939) (Jenett et al., 2012); (8) *R24H08-GAL4* (BDSC:49100) (Jenett et al., 2012); (9) *R25D01-GAL4* (BDSC:49122) (Jenett et al., 2012); (10) *repo-GAL4* (gift from Y-H. Sun); (11) *VT41043-GAL4* (gift from C-L. Wu); (12) *nSyb-GAL4* (Lin et al., 2012); (13) *UAS-DenMark,UAS-Dsyd-1::GFP* (Owald et al., 2015); (14) *UAS-mCD8::GFP*(LL4) (Lee and Luo, 1999); (15) *UAS-mCD8::GFP*(II) (Lee and Luo, 1999); (16) *UAS-mCD8::GFP*(III) (Lee and Luo, 1999); (17) *UAS-(FRT stop)myr-SmGdp-HA, UAS-(FRT stop)myr-SmGdp-V5, UAS-(FRT stop)myr-SmGdp-FLAG* (BDSC:64085) (Nern et al., 2015); (18) *MB247-LexA::p65, LexAop-rCD2::RFP* (Awasaki et al., 2011); (19) *LexAop-rCD2::RFP*(II) (Awasaki et al., 2011); (20) *UAS-hid, UAS-rpr* (gift from S. Waddell); (21) UAS-Dicer (Vienna Drosophila RNAi Center, VDRC:36148); (22) *20XUAS-FLPG5* (BDSC:55807) (Nern et al., 2011); (23) *R53F03-LexA::p65* (BDSC:53502) (Pfeiffer et al., 2010); (24) *R20G03-LexA::p65* (BDSC:52570) (Pfeiffer et al., 2010); (25) *R25D01-LexA::p65* (BDSC:53519) (Pfeiffer et al., 2010); (26) *R12C11-LexA::p65* (BDSC:52444) (Pfeiffer et al., 2010); (27) *hs-FLPG5* (BDSC:77140) (Nern et al., 2011); (28) *UAS-sema1a* (BDSC:65734) (Jeong et al., 2012); (29) *UAS-sema1.mEC-5xmyc* (BDSC:65739) (Jeong et al., 2012); (30) *UAS-sema1.mEC.Fc-5xmyc* (BDSC:65740) (Jeong et al., 2012); (31) *UAS-sema1a-RNAi-1* (BDSC:34320) (Perkins et al., 2015); (32) *UAS-sema1a-RNAi-2* (VDRC:36148) (Dietzl et al., 2007); (33) *UAS-plexA-RNAi* (BDSC:30483) (Pecot et al., 2013; Perkins et al., 2015); (34) *w, ala^E13^* (BDSC:9463) (Pascual and Préat, 2001); (35) *plexA^EY16548^* (BDSC:23097) (Bellen et al., 2011; Shin and DiAntonio, 2011); (36) *40A, sema1a^k13702^* (Kyoto Stock Center:111328) (Yu et al., 1998); (37) *Sema2b^f02042^, sema2a^p2^, UAS-mCD8::GFP/CyO* (gift from L. Luo) (Sweeney et al., 2011); (38) *hs-FLP122, 40A, tubp-GAL80* (gift from H-H. Yu); and (39) *w; sema1a Artificial exon/CyO; LexAop-tdTomato* (gift from L. Zipursky) (Pecot et al., 2013). The genotypes of the flies used in each figure are shown in **Table S2**. All flies were reared at 23 °C with 60% relative humidity, under a 12 h light: 12 h dark cycle. All our data were derived from male flies.

### Mosaic analysis with a repressible marker (MARCM) and Flip-out

Flies with proper genotypes were heat-shocked at 37 °C at different developmental stages for different durations (see **Table S2** for each of these parameters used in each figure). For flip-out clones induced at the adult stage, we waited for at least 3 days after heat-shock to ensure proper expression of the marker proteins. Brains were then dissected before performing immunofluorescence staining.

### Immunofluorescence staining

Fly brains were dissected in phosphate buffered saline (PBS; Sigma, P4417) and fixed in PBS containing 4% formaldehyde (Sigma, F8775) at room temperature for 20 min. The brains were then washed three times with PBS containing 0.5% Triton X-100 (PBST, Sigma-Aldrich, T8787); each time, the brains were incubated in PBST at room temperature for at least 20 min. After that, the brains were incubated in PBST containing 5% normal goat serum (blocking solution; normal goat serum, Jackson ImmunoResearch, 005-000-121) for 30 min at room temperature. After blocking, the brains were incubated overnight in blocking solution containing primary antibodies at 4 °C. The next day, the brains were washed three times, for 20 min each time, with PBST at room temperature before they were incubated overnight in blocking solution containing secondary antibodies at 4 °C. The next day, the brains were washed again three times, for 20 min each time, with PBST at room temperature. Then the brains were mounted with Gold Antifade reagent (Thermo Fisher Scientific, S36937). Antibodies used in each figure are listed in **Table S2**. In some cases, we performed sequential staining in which the brains were stained with the first set of primary and secondary antibodies before being incubated with the second set of primary and secondary antibodies. The staining procedures (single vs. sequential) for each figure are provided in **Table S2**. The primary antibodies used in this study are Rat anti-mCD8 (1:100; Invitrogen, MCD0800), Chicken anti-GFP (1:1000; Invitrogen, A10262), Rabbit anti-Dsred (1:100; Takara, 632496), Mouse anti-Trio (1:100; Developmental Studies Hybridoma Bank, DSHB, 9.4A), Mouse anti-FasII (1:100; DSHB, 1D4), Mouse anti-DLG (1:100; DSHB, 4F3), Mouse anti-TH (1:1000; Immunostar, 22941), Rat anti-FLAG (1:200; Novus Biologicals, NBP1-06721), Mouse anti-V5 (1:500; Bio-RAD, MCA1360), and Rabbit anti-V5 (1:500; Abcam, AB9116). The secondary antibodies used in this study are Goat anti-rat-488 (1:400; Jackson ImmunoResearch, 112-545-167), Goat anti-rabbit-Cy3 (1:400; Jackson ImmunoResearch, 111-165-144), Donkey anti-chicken-488 (1:400; Jackson ImmunoResearch, 703-545-155), Goat anti-mouse-Cy5 (1:400; Jackson ImmunoResearch, 115-175-166), Goat anti-mouse-Cy3 (1:400; Jackson ImmunoResearch, 115-165-166), Goat anti-mouse-488 (1:400; Invitrogen, A11029), and Goat anti-rabbit-Cy5 (1:400; Jackson ImmunoResearch, 111-175-144). The stained brains were imaged using a confocal microscope (Zeiss LSM880) and analyzed using Fiji/ImageJ(Schindelin et al., 2012) or Imaris x64 (v9.2.1, Oxford Instruments).

### Statistics

Statistical analysis was performed in Prism 8. All data were tested for normality using the Shapiro-Wilk normality test. No data points were excluded from the analysis. Since all our data are normally distributed, we used two-tailed unpaired Student’s *t-*test with Welch’s correction.

## Author contributions

C-H.L. and S.L. conceived the project and designed the experiments. C-H.L. performed the experiments and analyzed the data. S.L. wrote the paper and directed the research.

## Acknowledgments

We thank Y. Aso (Janelia Farm Research Campus, USA), G. Rubin (Janelia Farm Research Campus, USA), T. Lee (Janelia Farm Research Campus, USA), L. Luo (Stanford University, USA), Y-H. Sun (Academia Sinica, Taiwan), S. Waddell (University of Oxford, UK), H-H. Yu (Academia Sinica, Taiwan), C-L. Wu (Chang Gung University, Taiwan), L. Zipurski (University of California, Los Angeles, USA), and FlyLight (Janelia Farm Research Campus, USA), the Bloomington Drosophila Stock Center, Vienna Drosophila RNAi Center, Harvard TRiP RNAi Stock Center, Kyoto Stock Center, and Taiwan Fly Core for fly stocks. S. L. is funded by the Ministry of Science and Technology, Taiwan (107-2311-B-001-042-MY3) and Academia Sinica, Taiwan.

## Supplementary materials

**Figure S1.**
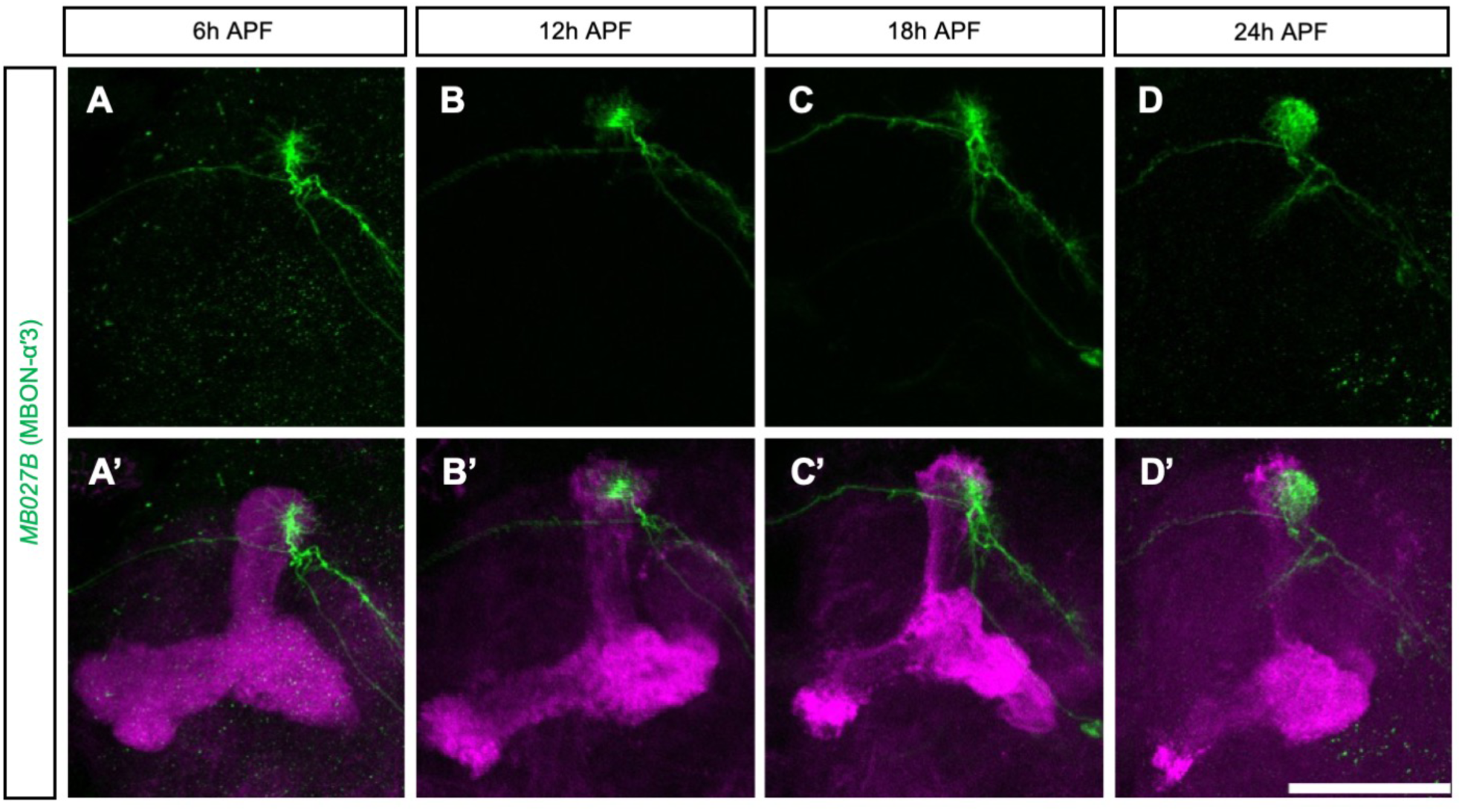
Innervation timings for MBON-αʹ3. **(A-D’)** Dendritic targeting of MBON-αʹ3 flip-out clones labeled with *UAS-mCD8::GFP* driven by *MB027B-splitGAL4* (green) at different hours after puparium formation (APF). All KC axons were labeled with *LexAop-rCD2::RFP* driven by *MB247-LexA::p65* (magenta, A’-D’). Scale bars: 50 µm. Detailed genotypes are listed in Table S2.

**Figure S2.**
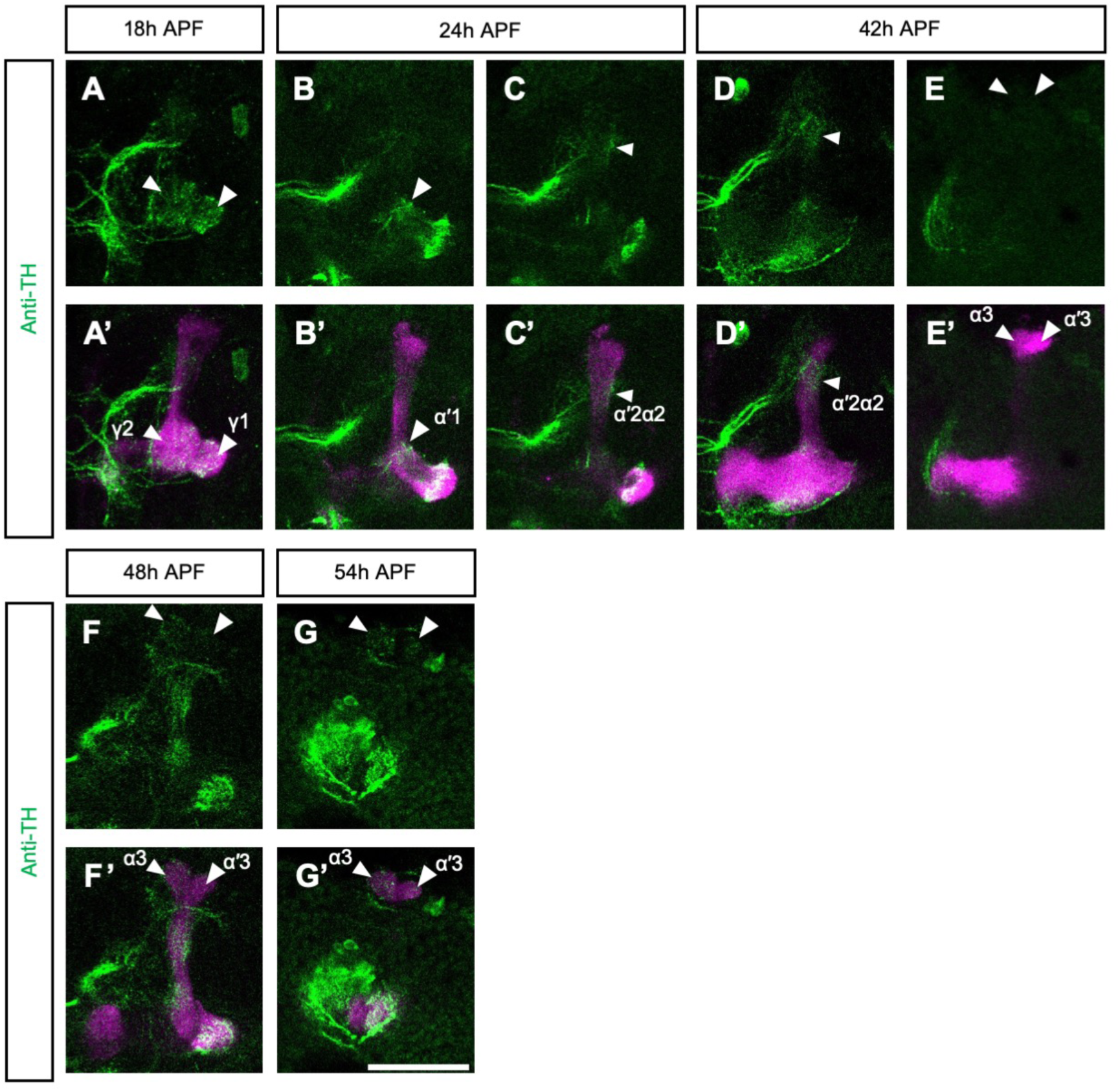
Anti-TH signals in the MB lobe during metamorphosis. Brains at different hours APF were stained with anti-TH antibody (green). All KC axons were labeled with *LexAop-rCD2::RFP* driven by *MB247-LexA::p65* (magenta). Note that TH signals in zones αʹ3 and α3 only start to appear at 48 h APF. Scale bars: 50 µm. Detailed genotypes are listed in Table S2.

**Figure S3.**
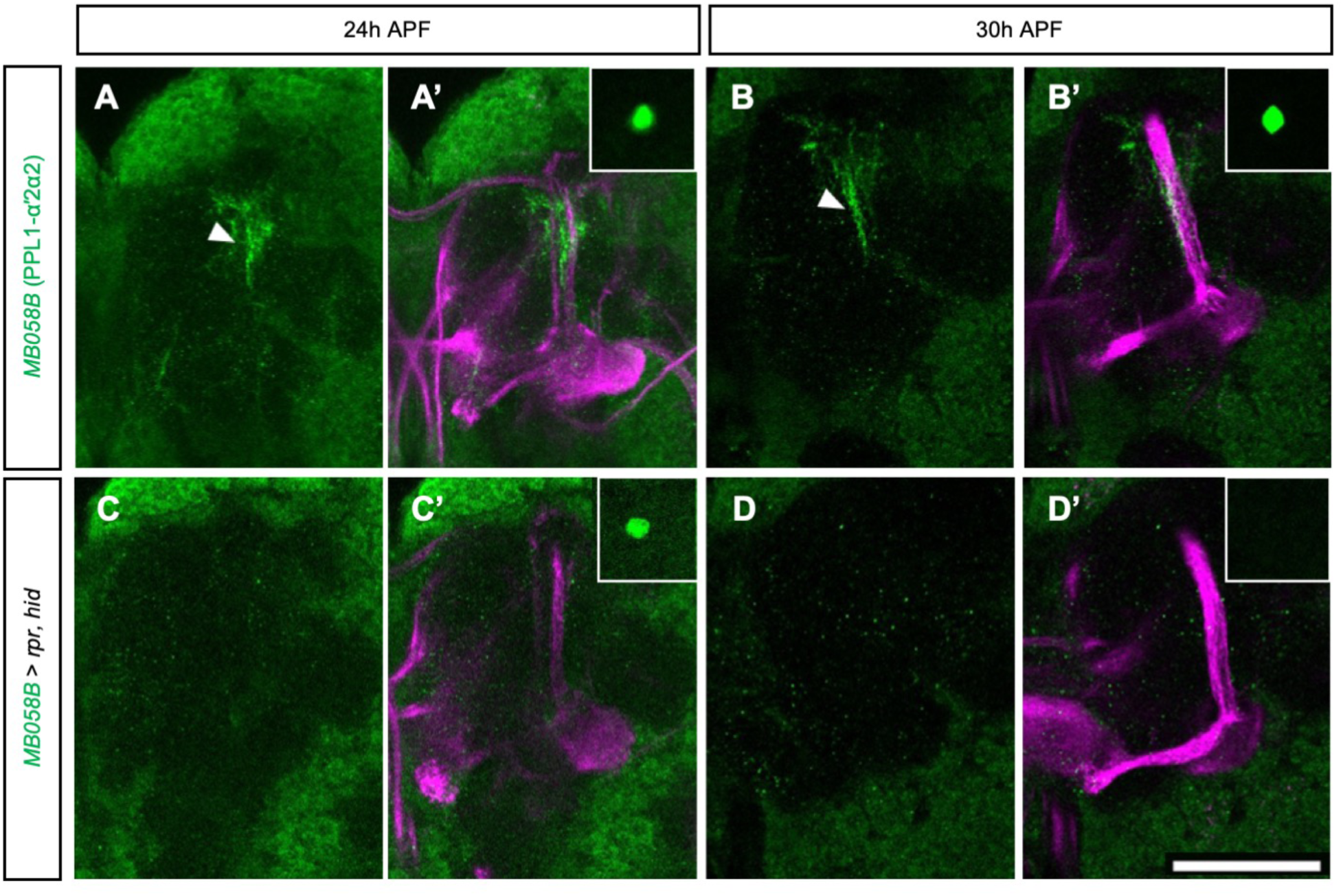
PPL1-αʹ2α2 DANs are ablated early in development. **(A-B’)** Axonal innervation patterns for PPL1-αʹ2α2 DANs labeled with *UAS-mCD8::GFP* driven by *MB058B-GAL4* (green) at 24 (A-A’) and 30 (B-B’) h APF. The brains were counterstained with anti-FasII antibody to label the α, β, and γ lobes (magenta, A’ and B’). Arrowheads indicate the innervations in the MB lobes (A and B). Insets in (A’ and B’) are the cell bodies of PPL1-αʹ2α2 DANs. **(C-D’)** As for (A-B’), except that *UAS-hid,rpr* was additionally expressed by *MB058B-GAL4* to ablate PPL1-αʹ2α2 DANs. Note that although the cell bodies of PPL1-αʹ2α2 DANs were still observed at 24 h APF, no axonal innervation in the lobes was apparent (C-C’). Cell bodies were completely ablated at 30 h APF (D-D’). Scale bars: 50 µm. Detailed genotypes are listed in Table S2.

**Figure S4.**
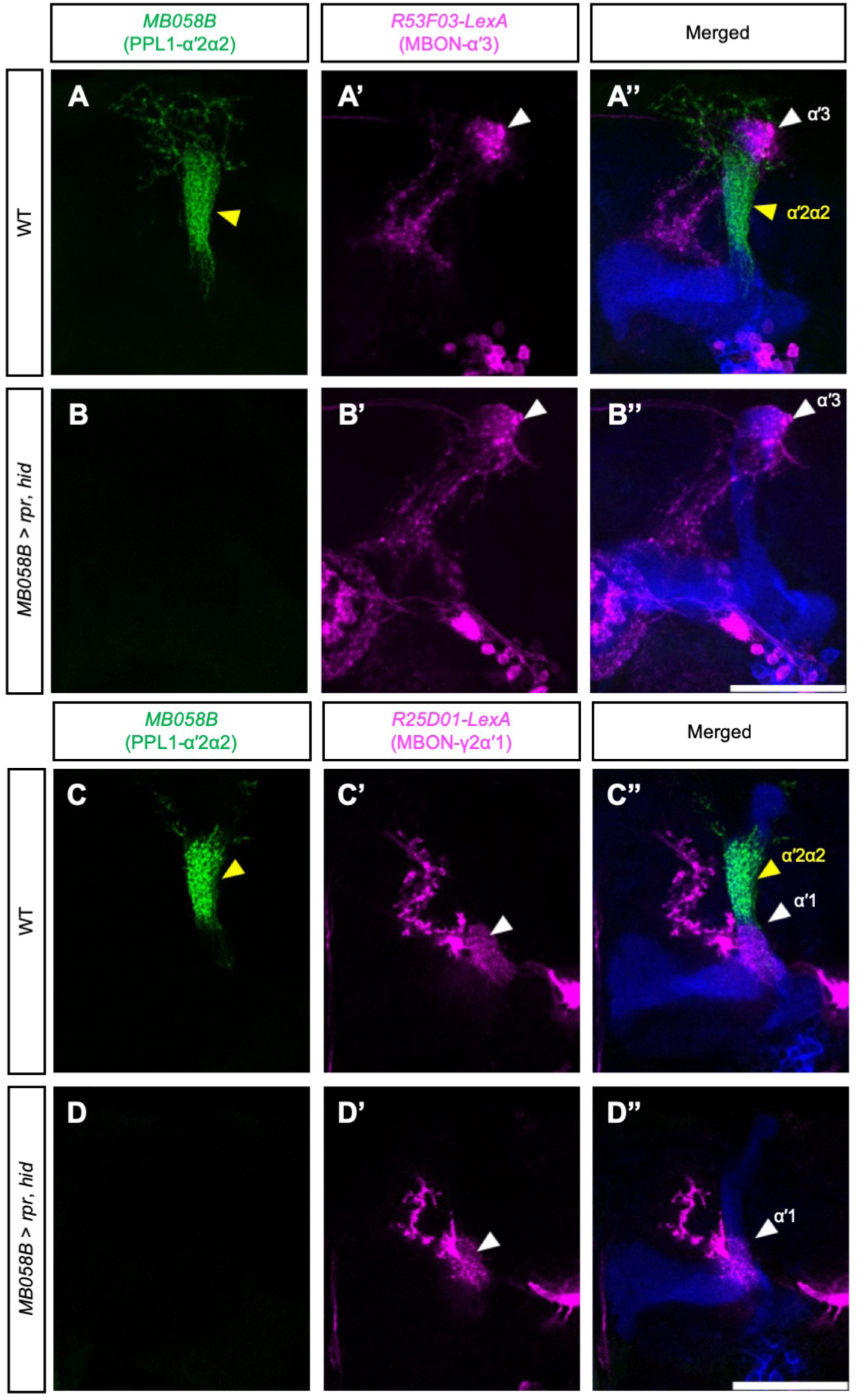
Ablation of PPL1-αʹ2α2 DANs has no effect on the dendritic targeting of neighboring MBONs. **(A-A’’)** A wild-type adult brain whose PPL1-αʹ2α2 DANs and MBON-αʹ3 were labeled, respectively, with *UAS-mCD8::GFP* driven by *MB058B-splitGAL4* (green, A and A’’) and *LexAop-rCD2::RFP* driven by *R53F03-LexA* (magenta, A’ and A’’). The brain was counterstained with anti-Trio antibody to label the αʹ, βʹ, and γ lobes (blue, A’’). The white arrowheads indicate the αʹ2α2 and αʹ3 innervations. **(B-B”)** As for (A-A’’), except that *UAS-rpr,hid* was additionally expressed using *MB058B-splitGAL4* to ablate PPL1-αʹ2α2 DANs. The white arrowheads indicate the innervation by MBON-αʹ3 dendrites (B’ and B’’). **(C-C’’)** As for (A-A’’), except that *R53F03-LexA* was replaced with *R25D01-LexA* to label MBON-γ2αʹ1 (magenta, C’-C’’). The white arrowheads indicate the αʹ2α2 and αʹ1 innervations. **(D-D’’)** As for (C-C’’), but *UAS-rpr,hid* was additionally expressed using *MB058B-splitGAL4* to ablate PPL1-αʹ2α2 DANs. The white arrowheads indicate the innervation by MBON-γ2αʹ1 dendrites of the αʹ1 zone (D’ and D’’). Scale bars: 50 µm. Detailed genotypes are listed in Table S2.

**Figure S5.**
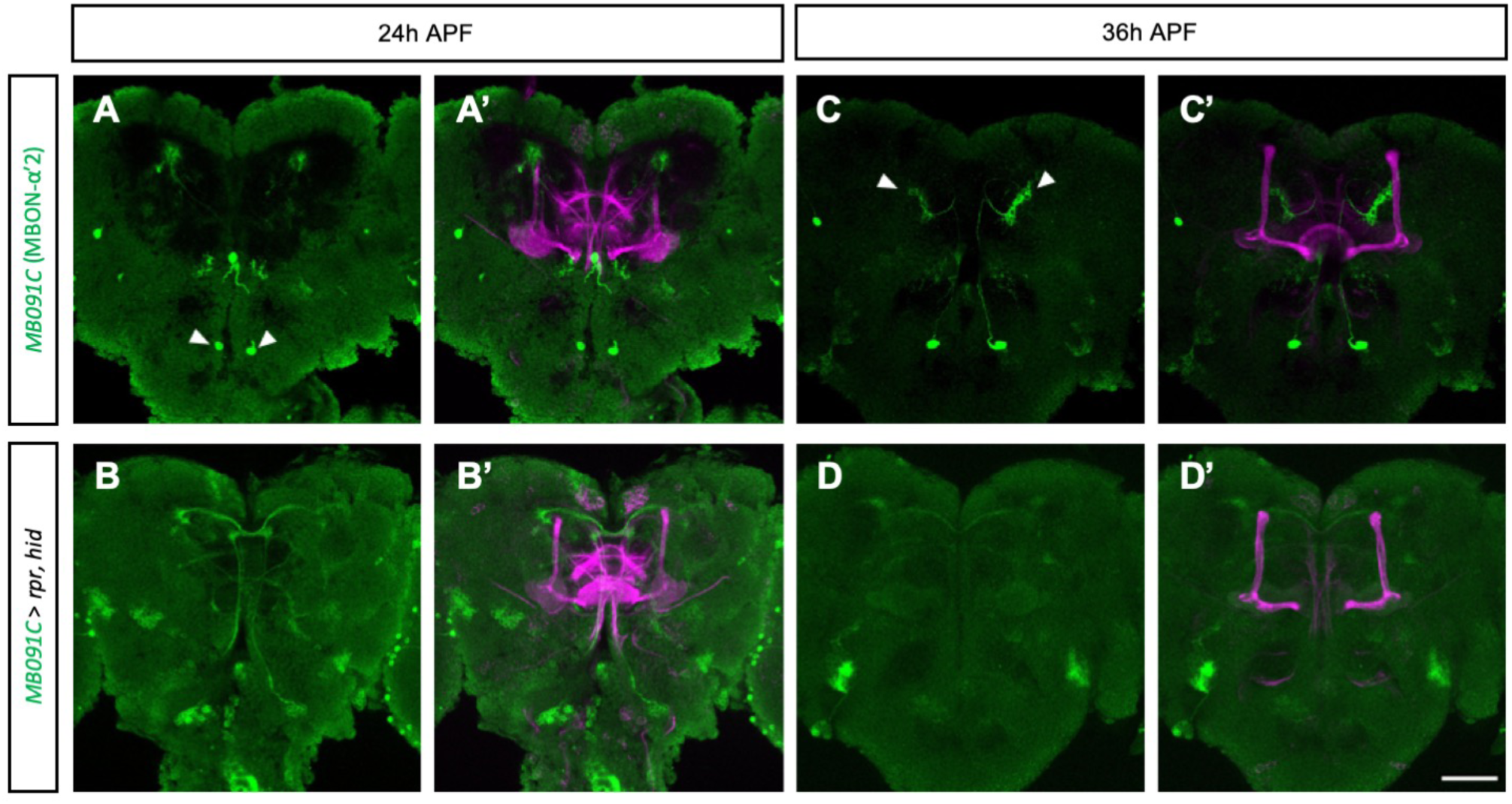
MBON-αʹ2 are ablated early in development. **(A-A’)** MBON-αʹ2 labeled with *UAS-mCD8::GFP* driven by *MB091C-splitGAL4* (green) at 24 h APF. The brains were counterstained with anti-FasII antibody to label the α, β, and γ lobes (magenta, A’). The white arrowheads in (A) indicate the cell bodies of MBON-αʹ2. **(B-B’)** As for (A-A’), except that *UAS-hid,rpr* was additionally expressed by *MB091C-splitGAL4* to ablate MBON-αʹ2. **(C-D’)** As for (A-B’), but the brain was dissected at 36 h APF. The arrowheads in (C) indicate the targeting of MBON-αʹ2 dendrites to the αʹ2 zones. Note that the cell bodies of MBON-αʹ2 were completely ablated as early as 24 h APF. Scale bars: 50 µm. Detailed genotypes are listed in Table S2.

**Figure S6.**
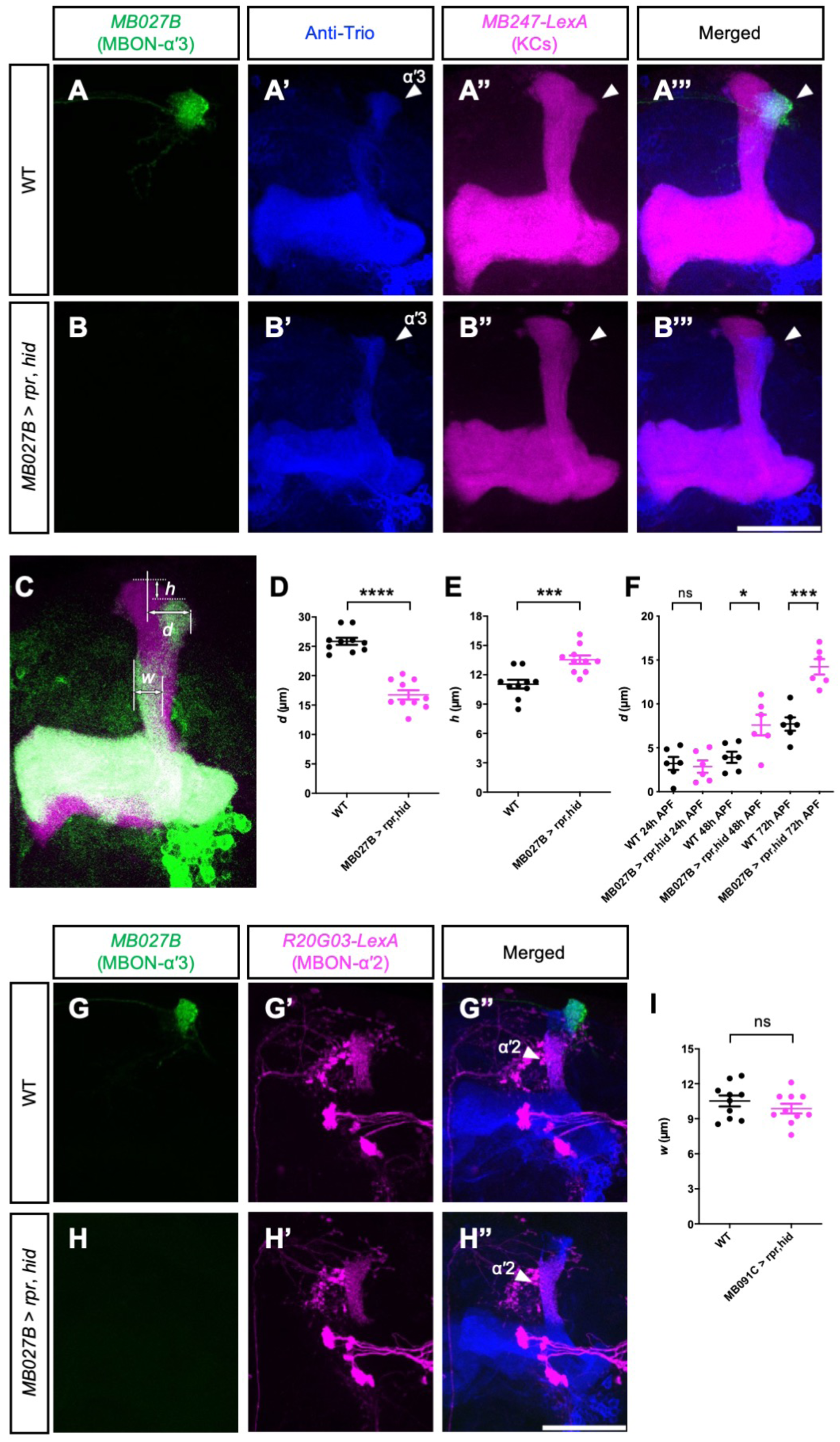
Ablation of MBON-αʹ3 impairs development of the αʹ lobe. **(A-A’’’)** A wild-type brain whose MBON-αʹ3 was labeled with *UAS-mCD8::GFP* driven by *MB027B-splitGAL4* (green, A and A’’’) and its KCs labeled with *LexAop-rCD2::RFP* driven by *MB247-LexA::p65* (magenta, A’’ and A’’’). The brain was also counterstained with anti-Trio antibody to label the αʹ, βʹ and γ lobes (blue, A’ and A’’’). The white arrowheads in (A’-A’’’) indicate the αʹ3 zone. **(B-B’’’)** As for (A-A’’’), except that MBON-αʹ3 was ablated with *UAS-rpr,hid* driven by *MB027B-splitGAL4*. **(C)** The parameters used to measure the position of the αʹ3 zone relative to the α lobe tip (*h* and *d*) and the width of the αʹ2 zone (*w*). *d* was measured as the horizontal distance between the most lateral position of the αʹ3 zone and the middle of the α lobe tip. *h* was measured as the vertical distance between the α and αʹ lobe tips. *w* was measured as the horizontal distance between two points, one on the inner border and the other on the outer border of the αʹ2 zone. The points were selected as the position closest to the brain midline on each border. **(D)** The lateral position of the αʹ3 zone (*d*) was reduced when MBON-αʹ3 was ablated (p<0.0001, n = 10, two-tailed unpaired *t-*test with Welch’s correction). **(E)** The relative height of the αʹ lobe was reduced (increased in *h*) when MBON-αʹ3 was ablated (p=0.0009, n = 10, two-tailed unpaired *t-*test with Welch’s correction). **(F)** Growth of the αʹ lobe was impaired when MBON-αʹ3 was ablated. Growth was measured as the relative positions of the tips of the α and αʹ lobes (*h*) at 24, 48, and 72 h APF (24 h APF: p=0.7224, n = 6; 48 h APF: p=0.0263, n = 6; 72 h APF: p=0.002, n = 6; two-tailed unpaired *t-*test with Welch’s correction). **(G-G’’)** A wild-type brain whose MBON-αʹ3 was labeled with *UAS-mCD8::GFP* driven by *MB027B-splitGAL4* (green, G and G’’) and its MBON-αʹ2 labeled with *LexAop-rCD2::RFP* driven by *R20G03-LexA* (magenta, G’ and G’’). The brain was also counterstained with anti-Trio antibody to label the αʹ, βʹ and γ lobes (blue, G’’). The white arrowheads in (G’’) indicate the αʹ2 zone. **(H-H’’)** As for (G-G’’), except that the MBON-αʹ3 was ablated using *UAS-rpr,hid* driven by *MB027B-splitGAL4*. **(I)** The width of the αʹ2 zone (*w*) was not altered upon MBON-αʹ2 dendrite ablation (p=0.3051, n = 10, two-tailed unpaired *t-*test with Welch’s correction). Scale bars: 50 µm. Detailed genotypes are listed in Table S2.

**Figure S7.**
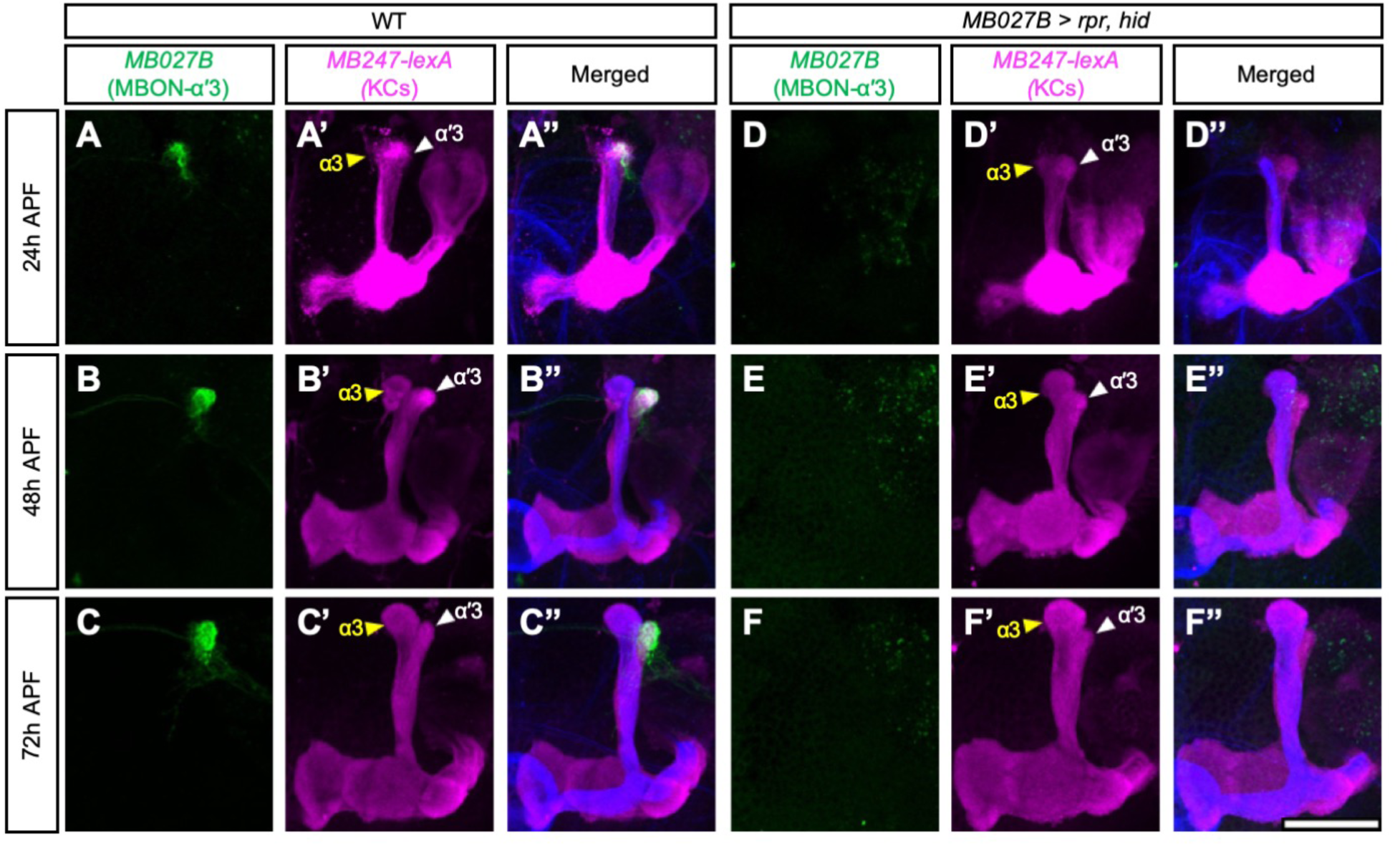
Development of the αʹ lobe without MBON-αʹ3. **(A-C’’)** MB lobes labeled with *LexAop-rCD2::RFP* driven by *MB247-LexA::p65* (magenta, A’-A’’, B’-B’’, and C’-C’’) at 24 (A-A’’), 48 (B-B’’), and 72 (C-C’’) h APF. MBON-αʹ3 was simultaneously labeled with *UAS-mCD8::GFP* driven by *MB027B-splitGAL4* (green; A, A’, B, B’, C and C’). The brains were counterstained with anti-FasII antibody to label the α, β, and γ lobes (blue; A’’, B’’ and C’’). Yellow and white arrowheads indicate the tips of the α and αʹ lobes, respectively. **(D-F’’)** As for (A-C’’), except that *UAS-hid,rpr* was additionally expressed by *MB027B-splitGAL4* to ablate MBON-αʹ3. Scale bars: 50 µm. Detailed genotypes are listed in Table S2.

**Figure S8.**
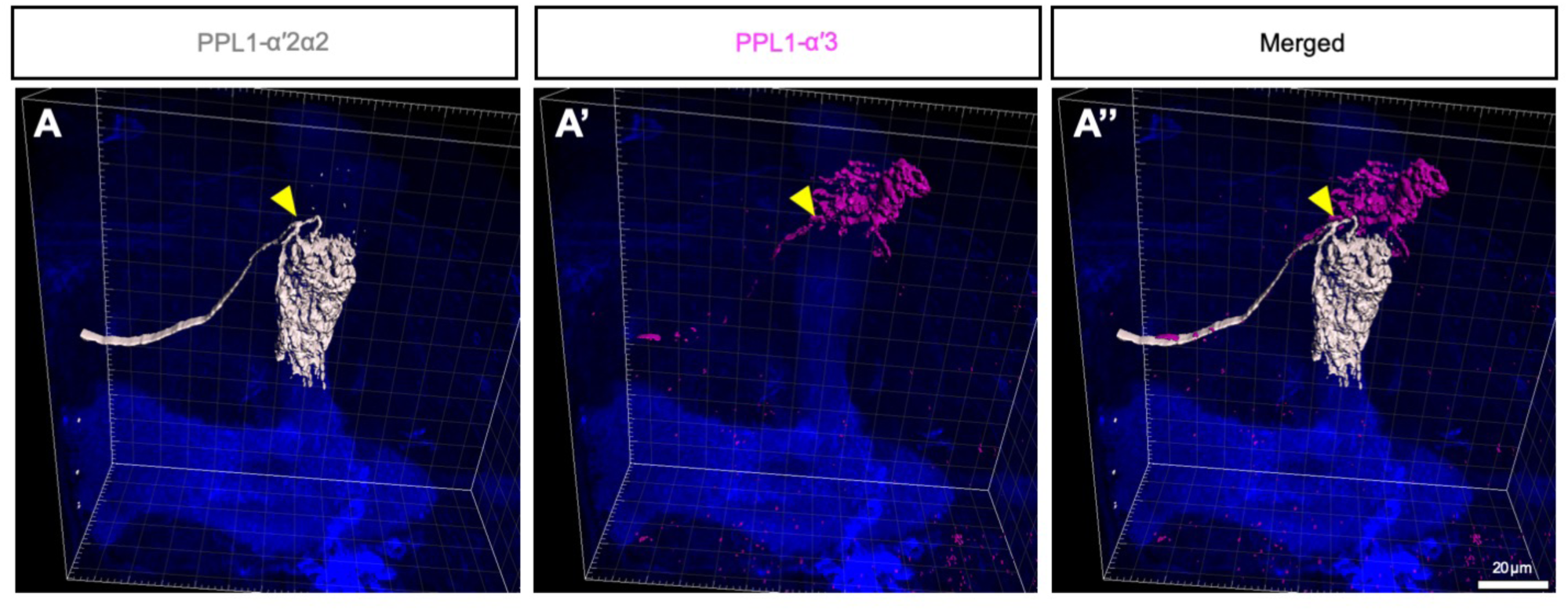
Multi-color flip-out clones of PPL1-αʹ2α2 and MBON-αʹ3 DANs in the same adult brain. **(A-A’’)** Multi-color flip-out clones of a PPL1-αʹ2α2 DAN (labeled by both V5-tag and FLAG-tag, white; A and A’’), and a PPL1-αʹ3 DAN (labeled by V5-tag, magenta; A’-A’’) in the same adult brain. Images were processed in Imaris. The yellow arrowheads indicate the common entry point in the MB lobes for both neuron types. Scale bars: 20 µm. Detailed genotypes are listed in Table S2.

**Figure S9.**
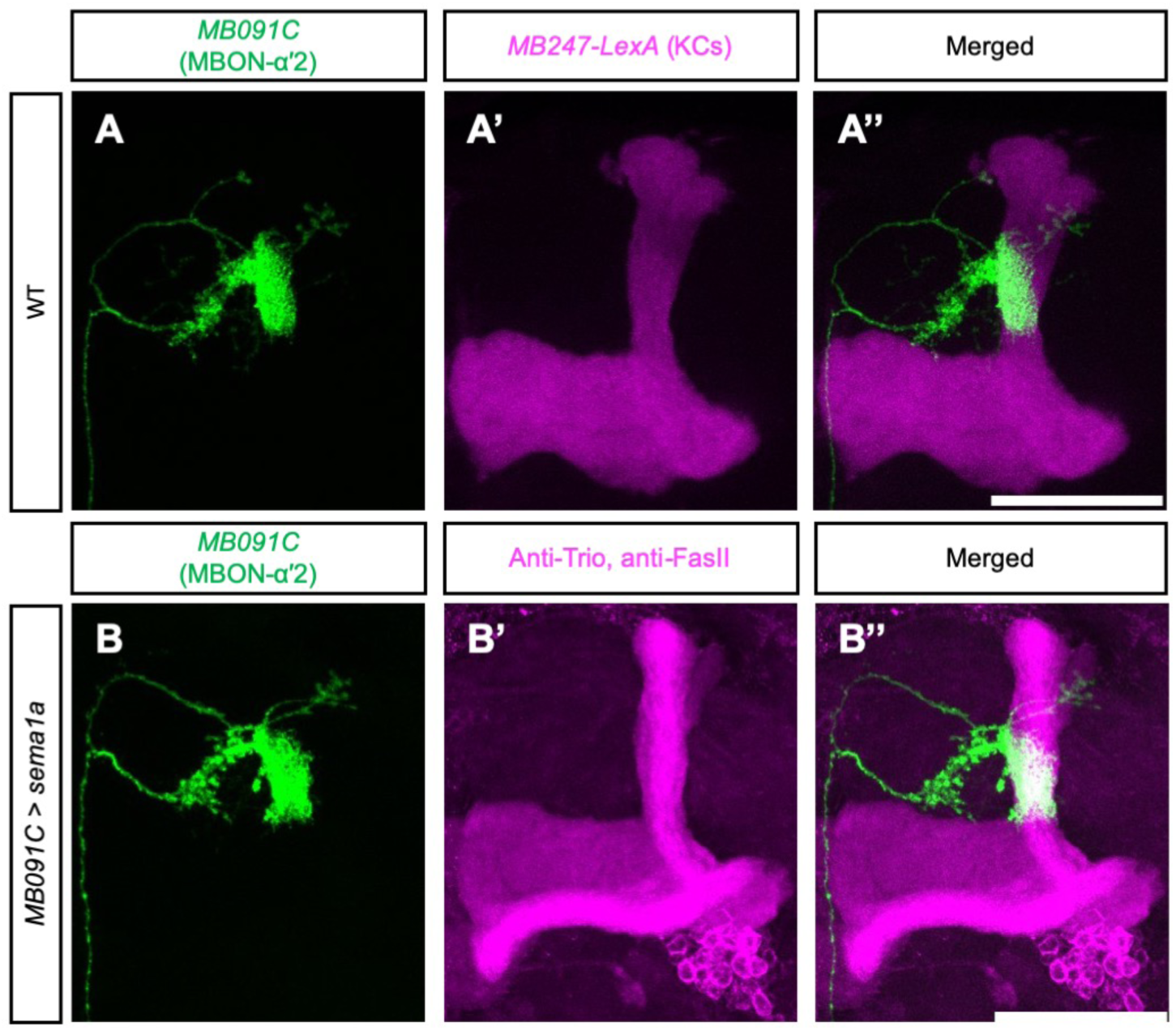
Targeting of MBON-αʹ2 dendrites is not affected by *sema1a* overexpression. **(A-A’’)** Targeting of MBON-αʹ2 dendrites labeled by *UAS-mCD8::GFP* driven by *MB091C-splitGAL4* (green, A and A’’). All KCs in (A-A’’) were labeled with *LexAop-rCD2::RFP* driven by *MB247-LexA::p65* (magenta, A’ and A’’). The brain in (B-B’’) was counterstained with anti-Trio and anti-FasII antibodies to label all MB lobes (magenta, B’ and B’’). **(B-B’’)** As for (A-A’’), except that *UAS-sema1a* was additionally expressed in MBON-αʹ2 by *MB091C-splitGAL4*. Scale bars: 50 µm. Detailed genotypes are listed in Table S2.

**Figure S10.**
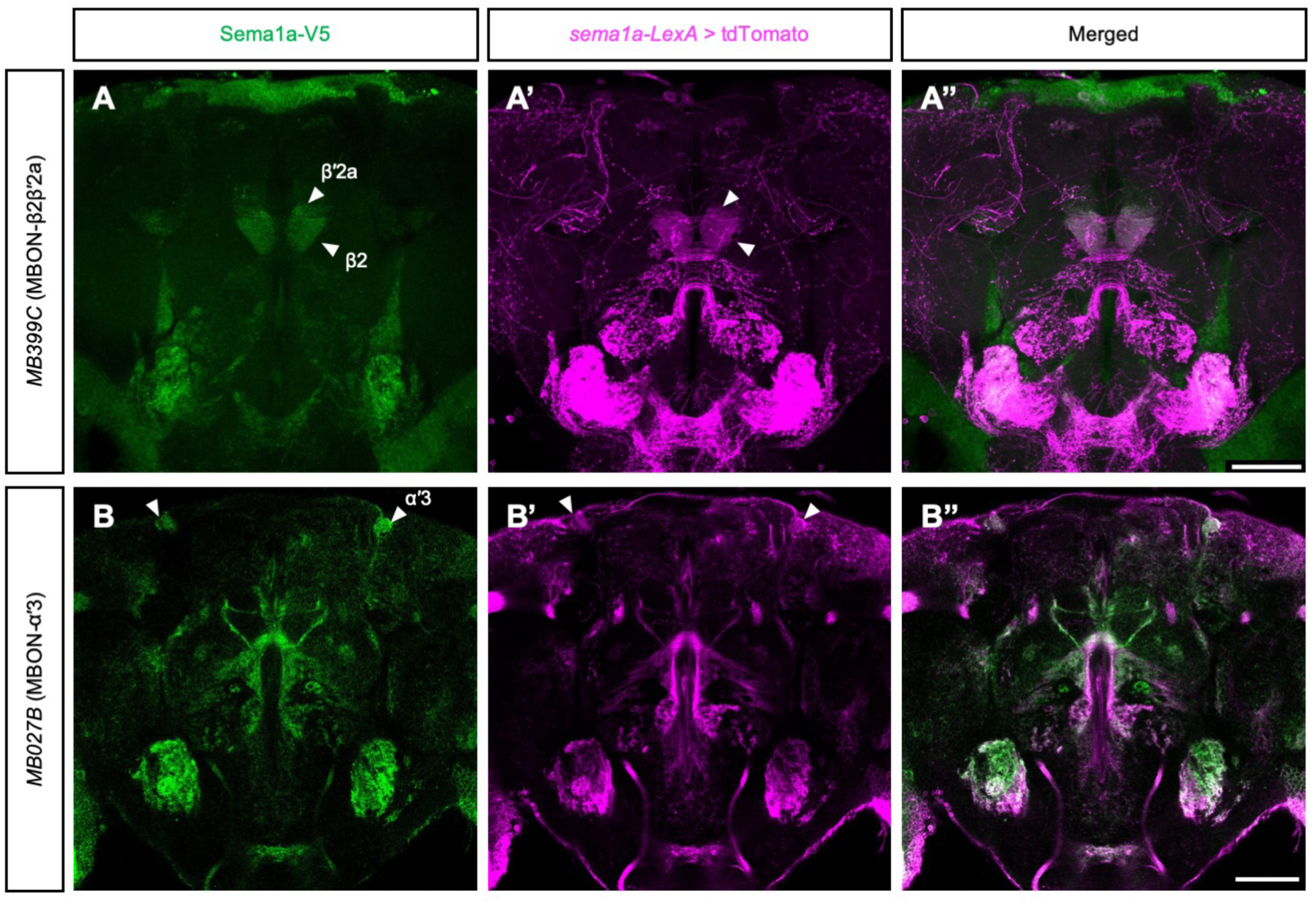
MBON-β2βʹ2a and MBON-αʹ3 express *sema1a*. **(A-B’’)** Expression of V5-tagged endogenous Sema1a (A and B) and tdTomato driven by the endogenous *sema1a* enhancer (A’ and B’) in MBON-β2βʹ2a (A-A’’) and MBON-αʹ3 (B-B’’) dendrites. Arrowheads in (A-A’) indicate the innervations by MBON-β2βʹ2a dendrites in the β2 and βʹ2a zones. Arrowheads in (B-B’) indicate the innervations of MBON-αʹ3 dendrites in the αʹ3 zones. *MB399C*-*splitGAL4* and *MB027B-splitGAL4* were used to remove the stop cassette from *sema1a* Artificial exon in MBON-β2βʹ2a and MBON-αʹ3, respectively. Scale bars: 50 µm. Detailed genotypes are listed in Table S2.

**Figure S11.**
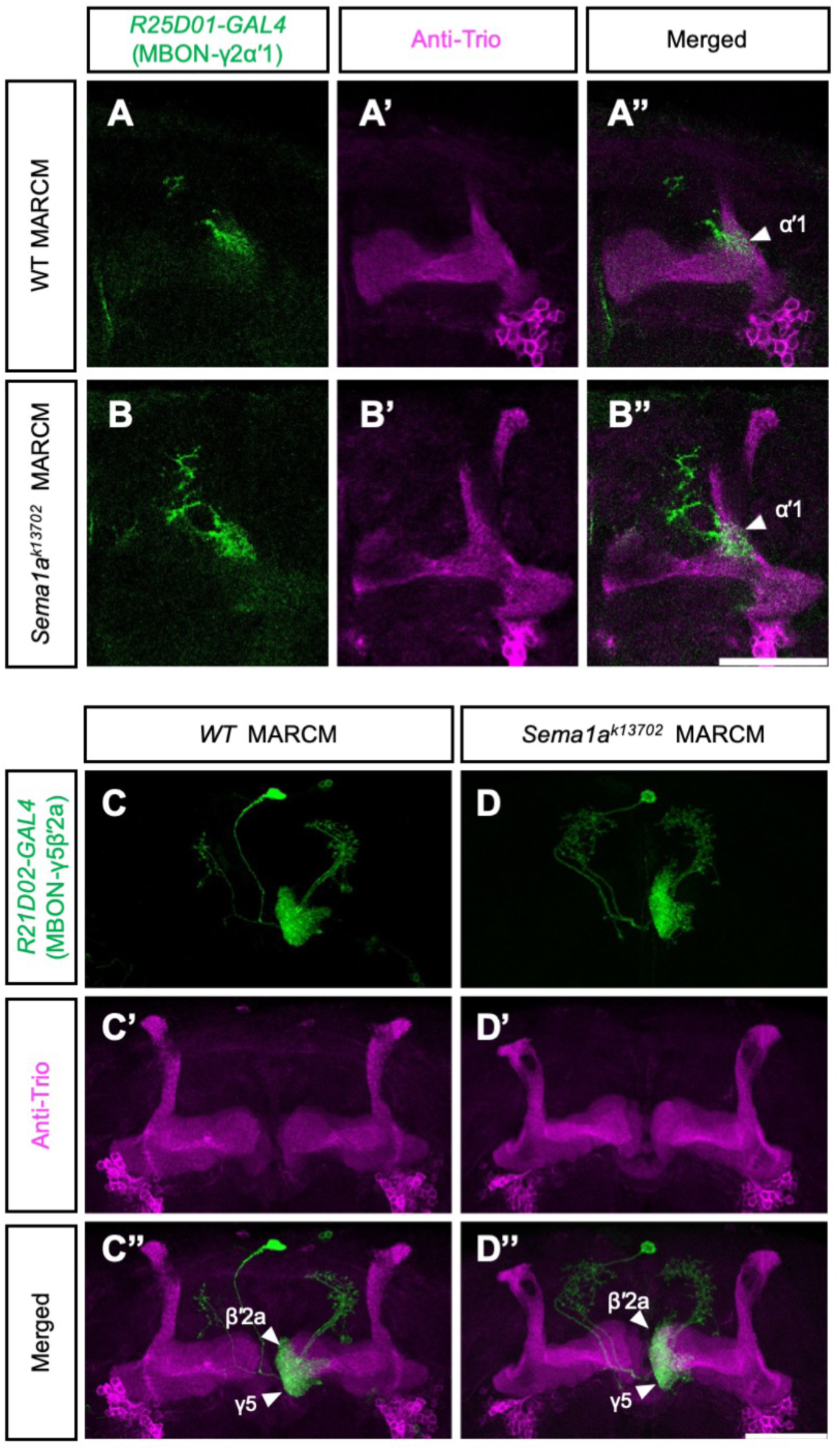
*sema1a* is not required for dendritic targeting of MBON-γ2αʹ1 and MBON-γ5βʹ2a. **(A-B’’)** Wild-type (A-A’’) or *sema1a^k13702^* (B-B’’) MARCM clones for MBON-γ2αʹ1 labeled with *UAS-mCD8::GFP* driven by *R25D01-GAL4* (green; A, A’’, B and B’’). The brains were counterstained with anti-Trio to label the αʹ, βʹ and γ lobes (magenta; A’-A’’ and B’-B’’). Arrowheads in (A’’ and B’’) indicate the αʹ1 zone. **(C-D’’)** As for (A-B’’), except that the clones are of MBON-γ5βʹ2a labeled by *R21D02-GAL4*. Arrowheads in (C’’ and D’’) indicate the βʹ2a and γ5 zones. Scale bars: 50 µm. Detailed genotypes are listed in Table S2.

**Figure S12.**
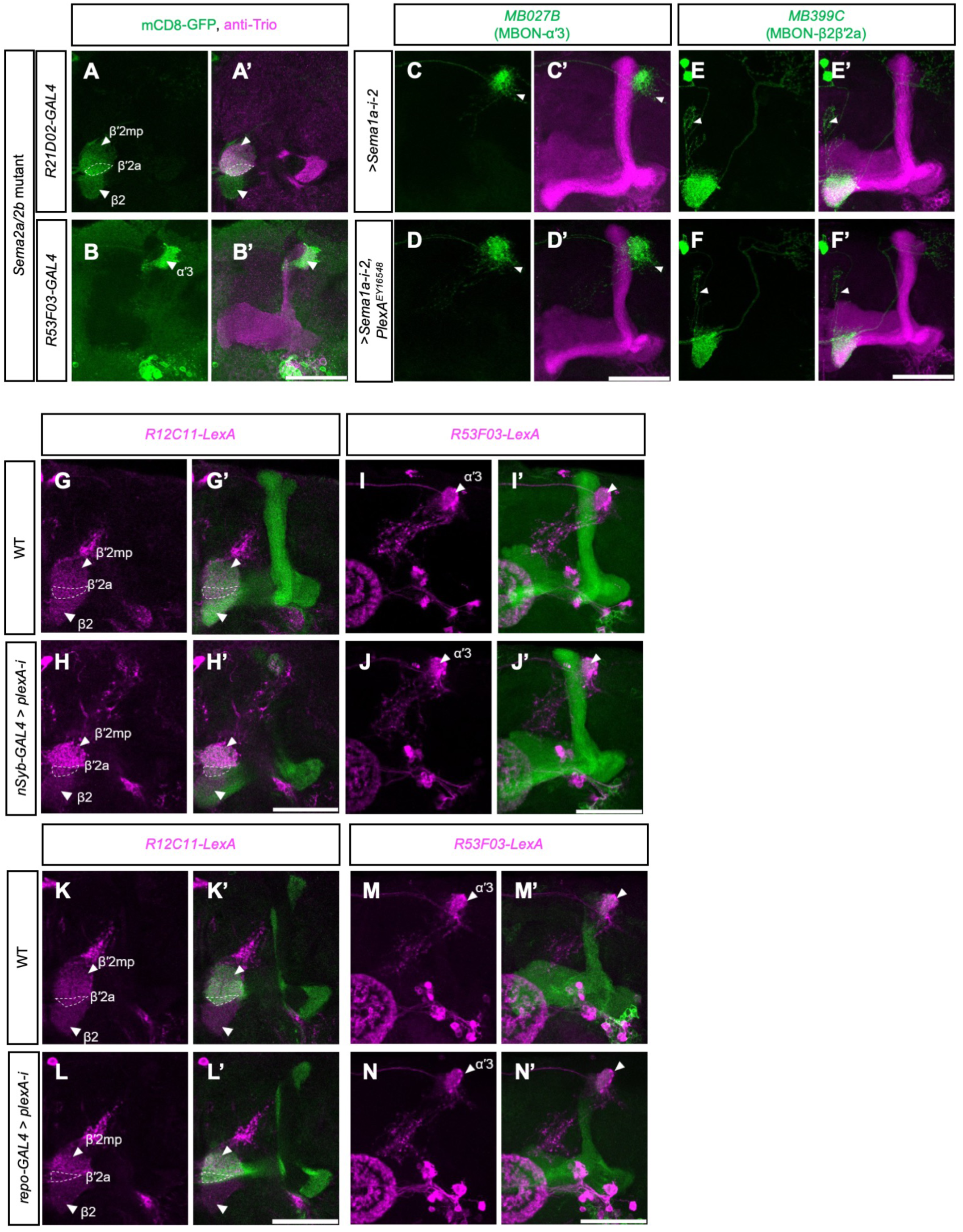
Canonical Sema1a ligands are not required for the dendritic targeting of MBONs. **(A-A’)** MBONs innervating the βʹ2mp, βʹ2a and β2 zones (indicated by white arrowheads and a dashed circle) were labeled with *UAS-mCD8::GFP* driven by *R21D02-GAL4* (green) in a *Sema2b^f02042^, sema2a^p2^* double mutant brain. The brain was counterstained with anti-Trio antibody to label the αʹ, βʹ, and γ lobes (magenta, A’). A signal focal plane is shown. Note that dendritic innervations in the MB lobe zones were largely unaffected. **(B-B’)** Dendritic innervation of MBON-αʹ3 labeled with *UAS-mCD8::GFP* driven by *R53F03-GAL4* (green) in a *Sema2b^f02042^, sema2a^p2^* double mutant brain. The brain was counterstained with anti-Trio antibody to label the αʹ, βʹ, and γ lobes (magenta, B’). Note that although the tip of the αʹ lobe is enlarged, the MBON-αʹ3 dendrites are largely confined to the αʹ3 zone (indicated by white arrowheads). **(C-D’)** MBON-αʹ3 dendrites expressing *UAS-sema1a-RNAi-2* and *UAS-mCD8::GFP* driven by *MB027B-splitGAL4* (green) in a wild-type (C-C’) or a *plexA^EY16548^* heterozygous mutant brain (D-D’). The brains were counterstained with anti-Trio and anti-FasII antibodies to label all MB lobes (magenta, C’ and D’). White arrowheads indicate the mistargeted dendrites. Note that the dendritic mistargeting phenotype was not exacerbated in the *plexA^EY16548^* heterozygous mutant brain (compare C and D as well as Fig. 8E). **(E-F’)** MBON-β2βʹ2a dendrites expressing *UAS-sema1a-RNAi-2* and *UAS-mCD8::GFP* driven by *MB0399C-splitGAL4* (green) in a wild-type (E-E’) or a *plexA^EY16548^* heterozygous mutant brain (F-F’). The brains were counterstained with anti-Trio and anti-FasII antibodies to label all MB lobes (magenta, E’ and F’). White arrowheads indicate the mistargeted dendrites. Note that the dendritic mistargeting phenotype was not exacerbated in the *plexA^EY16548^* heterozygous mutant brain (compare E and F as well as Fig. 8A). **(G-H’)** MBONs innervating the βʹ2mp, βʹ2a and β2 zones (indicated by white arrowheads and a dashed circle) were labeled with *LexAop-rCD2::RFP* driven by *R12C11-LexA* (magenta) in a wild-type brain (G-G’) or a brain in which *UAS-plexA-RNAi* was expressed pan-neuronally by *nSyb-GAL4* (H-H’). The brains were counterstained with anti-Trio and anti-FasII antibodies to label all MB lobes (green, G’and H’). **(I-J’)** MBON-αʹ3 waslabeled with *LexAop-rCD2::RFP* driven by *R53F03-LexA* (magenta) in a wild-type brain (I-I’) or a brain in which *UAS-plexA-RNAi* was expressed pan-neuronally by *nSyb-GAL4* (J-J’). The brains were counterstained with anti-Trio and anti-FasII antibodies to label all MB lobes (green, I’and J’). The white arrowheads indicate the αʹ3 zones. **(K-N’)** As for (G-J’), except that *UAS-plexA-RNAi* was expressed in all glia using *repo-GAL4* (L-L’ and N-N’). Scale bars: 50 µm. Detailed genotypes are listed in Table S2.

**Table S1. GAL4 lines examined for their potential to follow MBONs or DANs during their morphogenesis.**

**Table S2. Fly genotypes, immunostaining procedures, and heat-shock conditions for each figures.**

